# A rigidity phase transition of Stomatin condensates governs a switch from transport to mechanotransduction

**DOI:** 10.1101/2022.07.08.499356

**Authors:** Neus Sanfeliu-Cerdán, Borja Mateos, Carla Garcia-Cabau, Frederic Català-Castro, Maria Ribera, Iris Ruider, Montserrat Porta-de-la-Riva, Stefan Wieser, Xavier Salvatella, Michael Krieg

**Affiliations:** ICFO, Institut de Ciències Fotòniques, Castelldefels, Spain; IRB, Institute for Research in Biomedicine, Barcelona, Spain; ICREA, Barcelona, Spain

## Abstract

A large body of work suggests that biomolecular condensates ensuing from liquid-liquid phase separation mature into various material states. How this aging process is controlled and if the naive and mature phases can have differential functions is currently unknown. Using *Caenorhabditis elegans* as a model, we show that MEC-2 Stomatin undergoes a rigidity phase transition during maturation from fluid to viscoelastic, glass-like condensates that facilitate either transport or mechanotransduction. This switch is promoted by the SH3 domain of UNC-89/Titin/Obscurin through a direct interaction with MEC-2 and suggests a physiological role for a percolation transition in force transmission during body wall touch. Together, our data demonstrate a novel function for rigidity maturation during mechanotransduction and a previously unidentified role for Titin homologs in neurons.

## Main Text

The ability of cells to sustain and transmit mechanical force is intricately coupled to the controlled assembly, localization and mechanical properties of the constituent protein complexes (*1*). Stomatin family members are highly conserved scaffolding proteins that are involved in membrane organization (*2*) and modulate ion channel activity (*3-5*). The *Caenorhabditis elegans* (*C. elegans*) ortholog MEC-2 is essential for the sense of touch (*6,7*) and interacts with the N-terminus of the pore-forming subunit of the mechano-electrical transduction (MeT) channel MEC-4 at its stomatin domain (*8*). Like in other Stomatin proteins, the N and the C-termini have regions with low complexity composed of repetitive sequences of proline, glycines and serines and little homology to other proteins (fig. S1). In MEC-2, the C-terminal domain was hypothesized as a gating tether thought to modulate mechanosensitive ion channel open probability by a yet elusive protein-protein interaction (*9-11*).

Recent studies have shown that many proteins with intrinsically disordered domains have the propensity to separate from the bulk cytoplasm to form liquid-like condensates in a process akin to phase separation (*12-14*). Evidence is mounting that these liquid-like properties are impermanent and the condensates undergo a maturation to glass-like (*15*) and even solid aggregates (*16*). Such rigidity transitions have frequently been observed in proteins that form amyloid fibers in neurons and are implicated in disease, thus they are often hypothesized to drive neurodegeneration (*17, 18*). Accordingly, the processes leading to the condensation and fibril formation are of great interest in pathophysiology and drug discovery (*19*), especially the interactions within the condensate and their client proteins (*20*). *In vivo*, these condensates bind to, or coacervate other proteins, but their interaction are usually weak and transient in nature. It was proposed recently, that the transient cross-links evolve into stable interactions during a percolation transition (*19*), however, with deleterious consequences on their functionality. On the other hand, by definition, liquid-like condensates cannot sustain mechanical forces, and it is plausible that liquid-solid transitions also occur in physiologically relevant processes, for example in protein condensates at tight junctions (*21, 22*), focal adhesions (*23*) during mechanotransduction, or at scaffolding proteins found at sites of active mechano-electrical transduction.

### MEC-2 Stomatin exists as three phases *in vivo*

In order to shed light onto the molecular and cellular function of MEC-2 Stomatin *in vivo*, we first created transgenic animals carrying a single copy of fluorescently labeled MEC-2 and investigated its dynamics within touch receptor neurons (TRNs; movie S1) of immobilized animals. In agreement with previous results, we found that MEC-2 is distributed in discrete punctae along the sensory neurite (*8, 24*), however, with different dynamic signatures (Fig. 1A). MEC-2 resides in two distinct populations within the TRN neurite: in distal regions, punctae were predominantly static with little or no mobility. Closer to the cell body, however, we observed larger fluorescent punctae that rapidly trafficked towards the distal neurite with high processivity and occasional reversals, indicative for motor-driven, ballistic transport on short timescales (movie S1 and Fig. 1A). Based on their behavior and location, we termed the two pools mature and naive, respectively. Interestingly, the naive pool displays rich behavior reminiscent of liquid-like droplets (*14*) as we frequently observed fusion events between particles that encountered each other, fission of a single compartment and gel-like deformation presumably induced due to motor protein activity (Fig. 1B and movie S2). We thus reasoned that MEC-2 existed as phase separated biomolecular condensates (BMCs) *in vivo* with spatially distinct properties.

**Figure 1.**
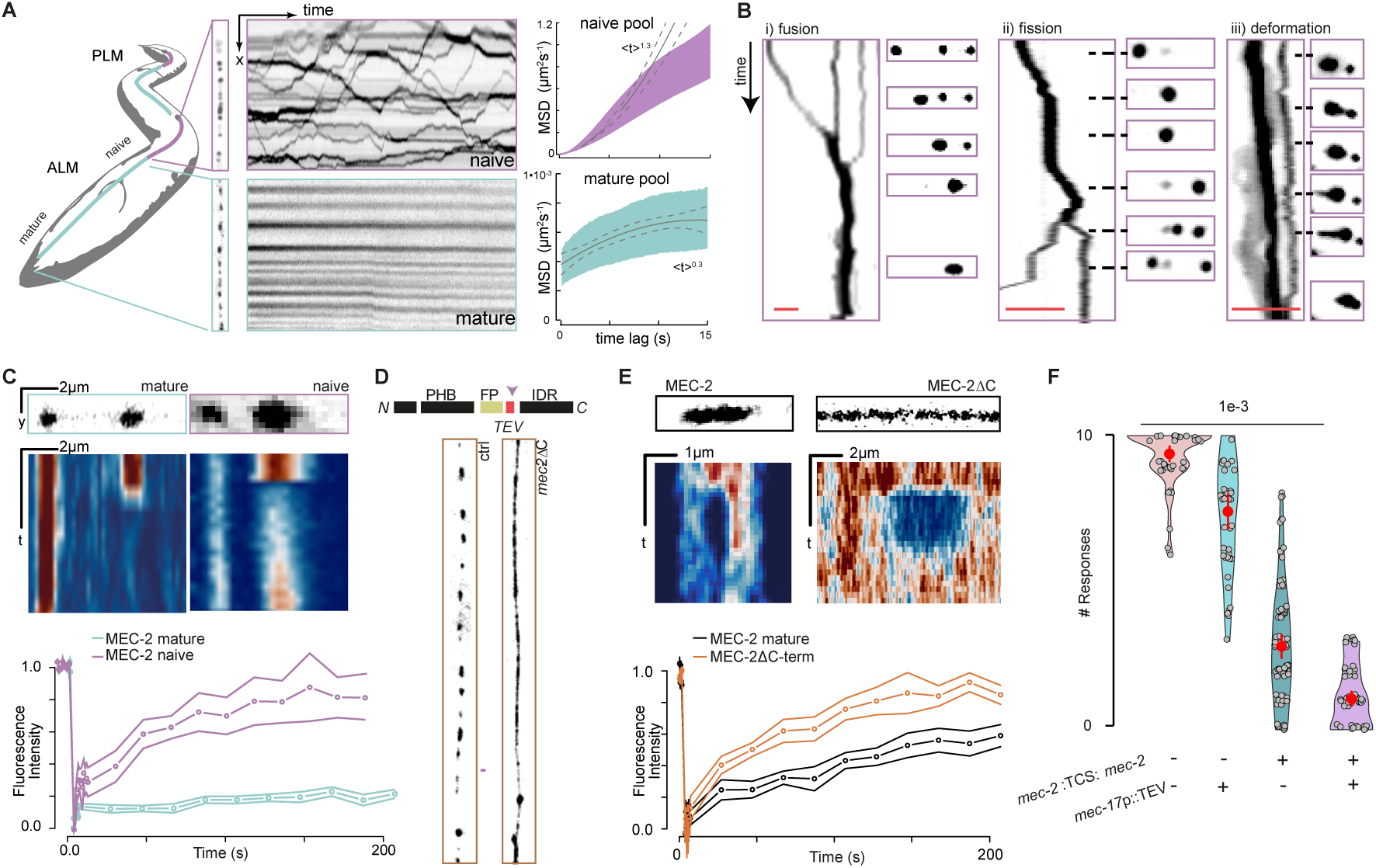
Dynamics of sol-gel transition. (**A**) Scheme of an animal indicating the naive and the mature pools of MEC-2 within an ALM touch receptor neuron. Representative image and kymograph of the naive, mobile pool and the mature, immobile pool of MEC-2 in the proximal and distal part of the TRN neurite, respectively. Mean square displacement (MSD) vs. timelag for MEC-2 in the naive and the mature fractions as indicated in the scheme to the left. The exponent of <*x*> *^n^* indicates the best fit to a power law. (**B**) Representative examples of MEC-2 condensates in the naive fraction of the TRN (labelled in purple in the scheme): (i) fusion, (ii) fission and (iii) deformation events plotted in a kymograph and snapshots during the course of the videos. Scale bar = 5 µm; sequence length = 5, 13 and 22 s, respectively. (**C**) Representative kymographs (top) and the fluorescence recovery dynamics after photobleaching of *mec-2*::mCherry in the mature pool of TRNs (green) and *mec-2*::mCherry in the naive pool expressed in hypodermal cells (purple). Mean±SD, N≥10 TRNs. (**D**) Schematic of the construct and representative images of MEC-2 distribution in TRNs of a *mec-2*::TSMod::TEV::*mec-2* animal (see scheme of the protein domains) in absence (left) and presence (right) of TEV protease. PHB, prohibitin domain; FP, fluorescent protein; TEV, Tobacco etch virus cleavage site; IDR, intrinsically disordered region. See quantification of interpunctum distances in fig. S5. (**E**) Representative kymographs (top) and the fluorescence recovery dynamics after photobleaching for fulllength MEC-2 (inside mature puncta, black) and conditionally truncated MEC-2 (orange). Mean±SD, N≥10 TRNs. (**F**) Violin plot of the body touch response derived from wildtype and conditionally truncted animals in absence or presence of TEV protease, and wildtype animals with only TEV protease expression. Circle indicates median, vertical bar indicates SD, N≥ 60 animals. *p*-value derived from Tukey HSD test.

To establish that MEC-2 forms different BMCs with distinct properties along the neurites *in vivo*, we compared the behavior of the mature and naive populations. Because the naive pool is very dynamic in TRNs, we established a transgenic model that expressed MEC-2 in hypodermal cells, where these condensates remained primarily static. We selectively bleached individual punctae in the mature and naive pools and imaged their fluorescence recovery after photobleaching during 3 minutes (FRAP; Fig. 1C). We observed that the naive, dynamic pool recovered quickly, while the mature, immobile pool did not recover at all during the course of the experiment. This suggests that the naive punctae are more liquid-like than the mature (D ≈ 0.2*µm*^2^/*s vs.* ≈ 0µ*m*^2^/*s*) presumably due to rapid exchange of MEC-2 between the condensed and the dilute phases (*25*).

We hypothesized that C-terminal domain drives the transition from the naive to the mature pool, which constitute the sites of mechano-electrical transduction. We engineered a fluorescent protein followed by a proteolytic TEV cleavage site into MEC-2, that separated the PHB domain and the C-terminus (Fig. 1D, arrow). In absence of the TEV protease, this construct localized in discrete regions within distal parts of the sensory neurite, indistinguishably from wildtype MEC-2 proteins. When we coexpressed the TEV protease under the TRN-specific *mec-17* promotor, we did not observe distinct punctae of the N-terminal, fluorescently labeled fragment, indicating that the C-terminal domain is necessary for condensation of MEC-2 into distinct domains (Fig. 1D). We next compared the dynamics of the continuous MEC-2 phase with the MEC-2 within the mature complexes in the distal neurite using FRAP. Whereas the cleaved MEC-2 lacking the C-terminal domain recovered quickly and completely after photobleaching (Fig. 1E and movie S3, D ≈ 0.25*µm*^2^/*s*), the MEC-2 inside the individual spots recovered only partially over the course of the experiments (D = 1·10*^−^*^3^*µm*^2^/*s*). To understand if the MEC-2 C-terminal domain and its propensity to form distinct punctae relate to its function in sensing mechanical touch, we assayed the effect of the TEV cleavage on aversive behavior. Consistent with the idea that the mature punctae constitute active sites of mechanotransduction, the conditional C-terminally truncated MEC-2 was not able to transduce external touch into avoidance behavior (Fig. 1F).

Taken together, MEC-2 forms distinct, liquid-like punctae that undergo a striking mobility and viscosity switch along the neurite, suggesting a role of the C-terminal domain for proper protein maturation, localization and function.

### MEC-2 forms condensates that mature into a viscoelastic, glass-like material

Protein structure prediction algorithms based on the AlphaFold2 (*26*) (Fig. 2A) did not fold the MEC-2 C-terminal domain in a well-defined three-dimensional structure. Indeed, this domain presents high propensity to be intrinsically disordered (ID) and undergo liquid liquid phase separation (LLPS) (*27*) (Fig. 2B, PScore (*28*)). To confirm this, we expressed and purified the domain (residues 371-481) and assessed its structural and phase separation properties *in vitro*. As expected in an ID protein, the solution-state nuclear magnetic resonance (NMR) ^1^H-^15^N spectrum displayed low ^1^H chemical shift dispersion (Fig. 2C) and the main chain chemical shifts largely corresponded to those of a statistical coil (Fig. 2D) (*29*). The intensities of the NMR signals across the sequence of the C-terminal domain where however non-uniform: in particular the signals corresponding to the region ^382^KKIRSCCLYKY^392^ appeared broad beyond detection (Fig. 2, C and D). This behavior has been observed in ID proteins that can undergo LLPS, in which low signal intensity identify regions of sequence involved in intra- or inter-molecular interactions that stabilize the phase separated state (*30, 31*). Next we tested the phase separation properties of the MEC-2 C-terminal domain *in vitro* and indeed observed that it forms liquid droplets (Fig. 2E and fig. S2A) upon heating (Fig. 2F) and at high ionic strength (fig. S2B), suggesting that the droplets are stabilized by hydrophobic interactions. The liquid droplets fused and their fluorescence quickly recovered after photobleaching, confirming their liquid character (Fig. 2, G and H, fig. S2C, and movie S4); these results are qualitatively equivalent to those observed for the BMCs formed by the full length protein *in vivo* (Fig. 1, B and C).

**Figure 2.**
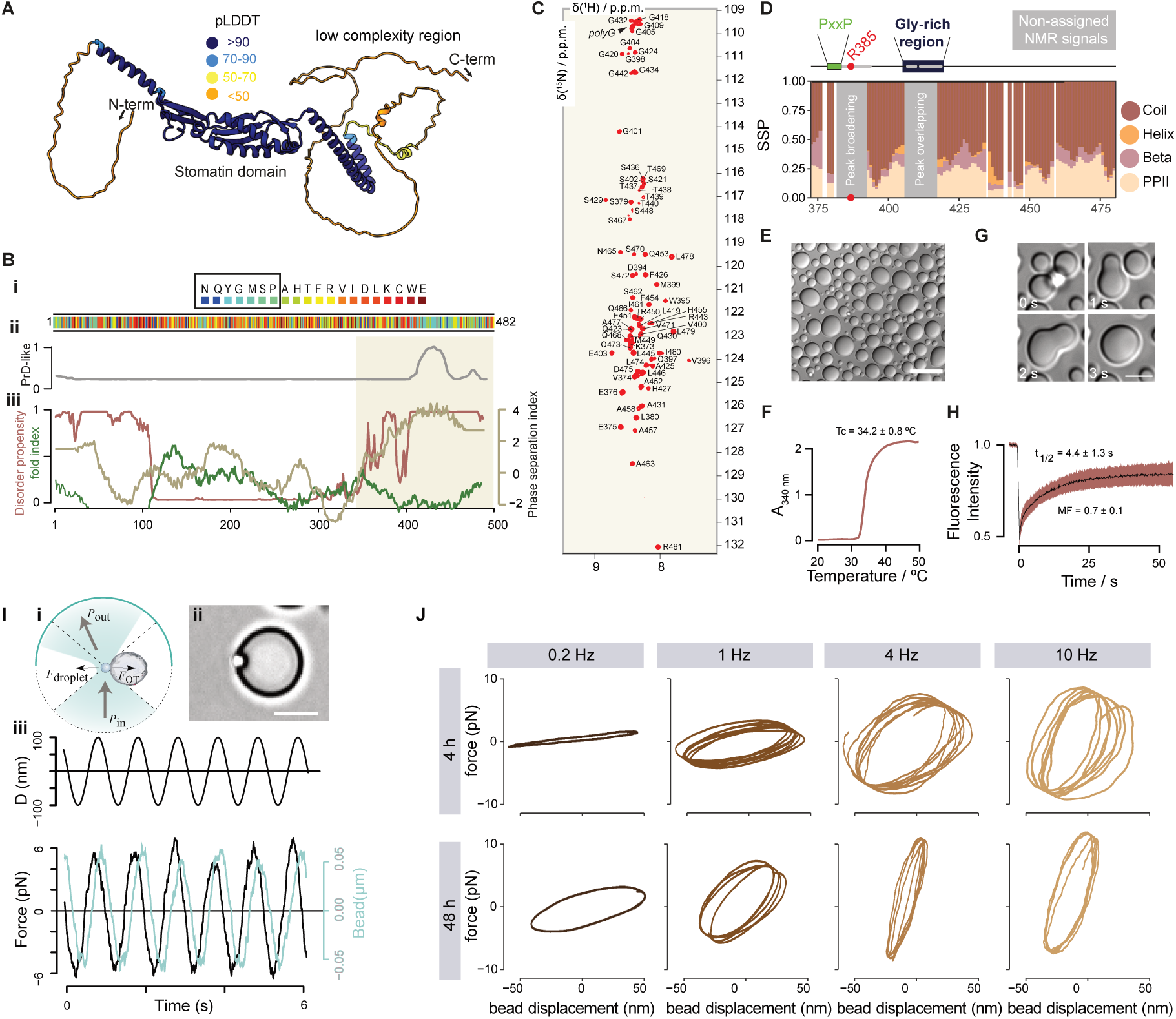
MEC-2 undergoes a viscoelastic switch during maturation from liquid to solid-like condensates. (**A**) AlphaFold prediction of MEC-2 Stomatin. Low confidence pLDDT values <50 suggest the presence of disorder in the N- and the C-terminal regions. (**B**) i) Color coded primary sequence of MEC-2 visualizes clustering of residues implicated in the formation of biomolecular condensates. ii) Prediction of prion-like sequences involved in higher order oligomerization or amyloid formation. iii) Bioinformatic analysis of the MEC-2 primary sequence according to its phase separation index (right axis, beige), disorder property calculation (left axis, red) and fold index (left axis, olive). (**C**) 2D NMR spectrum of MEC-2 C-terminal domain. (**D**) Secondary structure propensity of MEC-2 based on NMR chemical shifts. Schematic representation of MEC-2 motifs (PxxP, proline rich motif (green); and a glycine rich region (blue)). Grey boxes represent the non-assignable signals by NMR. (**E**) DIC microscopy image of MEC-2 liquid droplets *in vitro*, of a 400 µM sample with 2 M NaCl at 37 °C. Scale bar = 20 µm. (**F**) Apparent absorbance measurement as a function of temperature of 200 µM MEC-2 with 2 M NaCl. The indicated Tc value is the mean and standard deviation of three independent measurements. (**G**) MEC-2 droplets fusion *in vitro* at the indicated time points from movie S4. Scale bar = 5 µm. (**H**) FRAP experiment *in vitro* of 370 µM MEC-2 with 2 M NaCl at 20 °C. (**I**) i) Scheme of the optical tweezer based indentation assay, during which a trapped microsphere is oscillated into an immobilized droplet. ii) Representative picture showing the sphere in contact with the droplet. Scale bar = 5 µm. iii) Representative force-time signal of a typical sinusoidal rheology test. Upper graph indicates trap trajectory, lower graph the bead displacement and force. (**J**) Force-displacement plot (Lissajous-curve) showing the mechanical response of naive (4h) and mature (48h) droplet with increasing frequency. Opening of the circle indicates increased viscoelastic hysteresis.

Taken together, our data establishes that the MEC-2 C-terminal domain is intrinsically disordered and drives the formation of BMCs at distal sites of the TRN neurite, supporting the idea that the correct condensation and compartmentalization at the membrane is due to a liquid-solid transition.

We next directly tested the idea that MEC-2 undergoes spontaneous maturation from liquid to gel-like droplets. We set up an *in vitro* optical tweezer rheology assay (*15, 32*) to determine the viscoelastic properties of phase separated MEC-2 condensates (Fig. 2I). To do so, we performed step-indentations of 100 nm with a trapped microsphere and recorded the time-dependent viscoelastic stress relaxation (fig. S3, A and B). Freshly prepared droplets (t=0h) rapidly relaxed to a constant value close to the baseline level with a single time constant, indicating that naive MEC-2 condensates cannot store mechanical stress. In contrast, the relaxation time progressively increased with age for all time points tested (fig. S3C), reminiscent of a glass transition with an age-dependent increase in viscosity but without apparent effects on condensates stiffness (fig. S3, D and E). To better understand the mechanical behavior of MEC-2, we deformed the droplet sinusoidally (Fig. 2I iii) with an optically trapped microsphere at varying frequencies and recorded the resultant force (Fig. 2J). To our surprise, we observed that slow oscillations barely resulted in a measurable force for naive and mature droplets, but increased substantially for faster oscillations, especially for droplets of 48h in age (Fig. 2J). This suggests that the mechanical response of purified MEC-2 strongly depends on its age and on the rate of deformation, and might have pronounced consequences on the frequency selection during touch sensation *in vivo*.

Altogether, MEC-2 forms phase-separated liquid droplets that age into stress-storing, slowly relaxing condensates. We reasoned that the aging properties are important for mechanosensing.

### A C-terminal proline-rich motif is critical for mechanotransduction

Several MEC-2 point mutations have a strong touch defect without affecting trafficking of MEC-2 into the sensory neurite of the TRNs (*8*). One of these alleles, *u26*, encodes an arginine-to-histidine conversion at position 385 of its C-terminus (R385H) within the intrinsically disordered region (IDR), close to a proline rich motif (PRiM) with the consensus PxxP, reminiscent of SH3-binding domains (*33, 34*).

Animals carrying the R385H mutation neither displayed a behavioral response (fig. S4A) nor mechanoreceptor calcium transients in TRNs when punched into the body wall within a microfluidic chip (fig. S4, B and C, and movie S5) (*35*). Importantly, the MEC-2(R385H) localized to TRN dendrites indistinguishably to wildtype MEC-2 *in vivo* (fig. S5) and colocalized with the MeT MEC-4 channel (fig. S4D), indicating that they sort into the same biomolecular complex. In order to investigate the role of the C-terminus in an interaction in *trans*, we overexpressed a peptide derived from the MEC-2 C-terminus encompassing the wildtype or the mutant PRiM, and assayed the animals’s response to touch. Whereas the mutant PRiM did not interfere with touch when overexpressed in wildtype animals, the wildtype PRiM domain led to a significantly reduced touch response (fig. S4E). This indicates that the wildtype but not the mutant PRiM domain can competitively interfere with the sense of touch and raises the possibility that the MEC-2 PRiM acts in *trans*, through an interaction with an unknown binding partner, rather than misfolding.

To understand why a single residue completely abrogates a behavioral phenotype without any visible defect in trafficking and axonal localization, we expressed the R385H mutant MEC-2 C-terminal domain *in vitro* and studied its structural propensities and LLPS behaviour. We found that it formed droplets indistinguishable from wildtype MEC-2 (fig. S4, F-J). The NMR spectra of the R385H mutation, however, showed a strong signal intensity reduction close to the PRiM as compared to wildtype MEC-2 (fig. S4, K-M), indicating that the R385H point mutation affects MEC-2 homotypic interactions. Indeed, the mutant MEC-2 forms closer contacts within mature condensates *in vivo* compared to wildtype MEC-2, as determined through FRET experiments from doubly transgenic animals expressing MEC-2::Venus or MEC-2(R385H)::Venus and MEC-2::mCherry (fig. S4N), however, with unchanged viscosity (fig. S4 O-Q).

Together, these analyses suggest that the R385H mutant MEC-2 has stronger homotypical intermolecular interaction that involves a sticky region proximal to a PRiM SH3 binding motif and leads to a defective touch response.

### MEC-2/Stomatin and UNC-89/Titin/Obscurin co-assemble into punctate structures

One proposed function of BMCs is to locally concentrate binding partners to accelerate biochemical reactions (*14, 36*), e.g. nucleate actin filaments (*37*). Based on the stereotypic PRiM present in the C-terminal part, we hypothesized that MEC-2 interacts with an SH3 domain (*38*). There are in total 81 proteins with an SH3 domain in the *C. elegans* proteome (*39*), many of which are unrelated to a specifically neuronal function. In order to identify the potential interaction partners of the SH3 binding motif, we performed a neuronal RNAi feeding experiment (Fig. 3A, inset) to knock-down 35 of 41 proteins with an SH3-domain (*39*) that are reportedly expressed in TRNs (*40, 41*) (table S1) and available in the collection (*42, 43*). We used mutant animals for systemic RNAi (*44*), but specifically sensitized in TRNs. When we cultured these animals on bacteria expressing *mec-4* or *mec-2* RNAi constructs (*42, 45*), they significantly decreased their response to touch, whereas the empty vector did not affect touch sensitivity. Surprisingly, we found that only the knockdown of *unc-89*, a member of the Titin family (Fig. 3A and fig. S6A; (*46, 47*)), gave a reproducible and robust reduction in the response to touch. The incomplete knockdown was enhanced in the *unc-89*/*mec-2* double feeding, suggesting a role in the mechanotransduction pathway. Because the function of UNC-89 and Titin in general was previously unknown in neurons, we chose to study its role in neuronal physiology in general and touch in particular.

**Figure 3.**
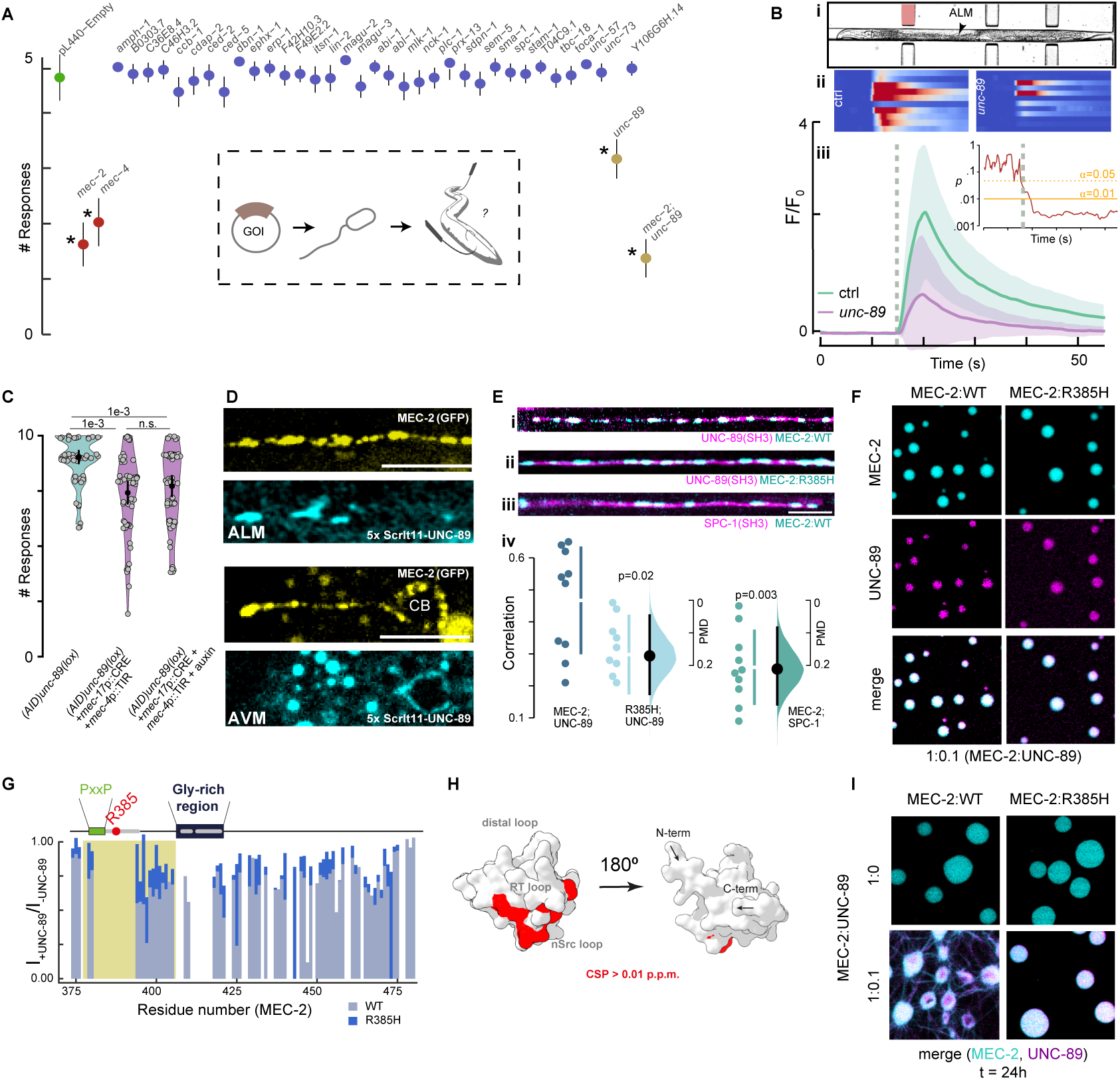
UNC-89/Titin/Obscurin is a component of the mechanoreceptor complex in TRNs. (**A**) Neuronal feeding RNAi screen with SH3-containing proteins naturally expressed in TRNs (see Methods and table S1). Mean±SD of head touch responses, N=40. Asterisk indicates a significant *p*-value compared to the empty vector (negative control) derived from nonparametric multiple comparison post-hoc Dunn’s test. *p*-values were: *mec-2*: 1e-19; *mec-4*: 3e-15; *unc-89*: 1e-7; *mec-2/unc-89*: 5e-37. Inset shows the scheme of the experimental pipeline. GOI, gene of interest. (**B, C**) UNC-89 knockout influence in *C. elegans* touch sensation. (**B**) (i) Brightfield image of an animal inside the body wall chip. (ii) Stacked kymographs of individual calcium recordings for ≥10 different animals. (iii) Average normalized fluorescence intensity ±SD for the GCaMP signal in a wildtype (green) or a UNC-89 KO animal (purple) upon body wall touch inside the microfluidic device (*35*). The calcium-independent mtagRFP-T as a control measurement is shown in fig S6F. Inset shows the p-value for each time point testing the Hypothesis H_0_: wt = *unc-89* with α=0.01. Pressure applied for 2 s at the time indicates as grey dotted line. (**C**) Violin plot showing the body touch response for TRN-specific knockout of *unc-89* by using double effect of CRE/loxP and auxin-induced degradation (AID) compared to the AID and loxP flanked control animals in absence of the *mec-17*p::CRE, the *mec-4*p::TIR or the auxin. Circle indicates mean, vertical bar indicates 95% confidence interval, N=60 animals. *p*-value derived from Tukey HSD test. (**D**) Representative images of split wrmScarlet(11)x5:*unc-89* complemented with TRN-specific *mec-4*p::wrmScarlet1-10 in *mec-2*::GFP background animals. Scale bar = 10 µm. CB, cell body. (**E**) Representative images of individual TRNs expressing a translational GFP fusion of the SH3 domain derived from UNC-89 and (i) MEC-2 wildype or (ii) MEC-2(R385H) mutant, and (iii) SPC-1 α-spectrin SH3 domain::GFP together with the wildtype MEC-2::mCherry. Scale bar = 10 µm. (iv) Altman-Gardner plot of the correlation between the SH3 domains and the MEC-2 condensates (*63*). (**F**) Confocal fluorescence microscopy images of 200 µM MEC-2 C-terminus (WT or R385H mutant) labeled with Alexa Fluor 647 together with UNC-89 SH3 domain labeled with Dy-Light 488, at a molar ratio of 1:0.1 (MEC-2:UNC-89), with 2 M NaCl at 37 °C. Scale bar = 20 µm. (**G**) Intensity ratio between the ^1^H-^15^N NMR spectra of the C-terminal domain of MEC-2 (WT (light blue) and R385H mutant (blue)) in the presence or the absence of the SH3 domain of UNC-89. (**H**) Structural map of the MEC-2 binding to the UNC-89 SH3 domain (represented as an AlphaFold model), derived from the chemical shift perturbations (CSP) in fig. S7D. Red residues correspond to CSP higher than 0.01 p.p.m (threshold). The binding is located at the surface between the RT and nSrc loops of the SH3, as previously described for other SH3 domains. (**I**) Confocal fluorescence microscopy images of 200 µM MEC-2 C-terminus (WT or R385H mutant) labeled with Alexa Fluor 647 without (1:0) or with (1:0.1) UNC-89 SH3 domain labeled with DyLight 488 (molar ratio MEC-2:UNC-89), with 2 M NaCl after 24 hours of sample incubation at 37 °C. Maturation into fibrilar structures was observed for MEC-2 (WT):UNC-89, but not for MEC-2(R385H):UNC-89 or in absence of UNC-89. Scale bar = 20 µm.

We first confirmed *unc-89* expression in TRNs with a driver construct that contained a GFP fusion to the genomic fragment including its promotor and the first three exons. In addition to TRNs, we observed noticeable expression in motor neurons and some neurons in the head and the tail, apart from the previously described expression in body wall muscles (fig. S6B and ref (*46*)). Next, we generated a knockout of the largest SH3-containing isoforms using CRISPR/Cas9 (fig. S6, C and D). Even though these knockout animals had a modest reduction in the behavioral response to touch compared to wildtype animals (fig. S6E), they consistently displayed lower calcium activity after mechanical stimulation in the body wall chip (*35*) as compared to controls (Fig. 3B and fig. S6F). To further confirm the cell-specific role of UNC-89, we conditionally deleted *unc-89* through the combination of site-specific CRE/lox recombination and auxin-induced degradation (AID, (*48*)). Coexpression of a panneuronal (fig. S6G) or a TRN-restricted CRE recombinase and TIR ligase after feeding the animals with auxin led to a slight but significantly decreased animals’s touch response (Fig. 3C). Collectively, this indicates that UNC-89/Titin/Obscurin is involved in the sense of touch in TRNs of *C. elegans*.

We next sought to decipher how UNC-89 enables full touch sensitivity in TRN neurites. Several components of the MeT channel complex associate into a punctate pattern (*8, 24*) and we asked if the *unc-89* knockout disrupts the punctate distribution of MEC-2 in the distal neurites. Even though the overall distribution of MEC-2 in the UNC-89 knockout is similar to the wildtype animals (fig. S5), we consistently observed that the median interpunctum interval (IPI) is significantly smaller (IPI_wt_ = 3.1µm vs IPI_u89_ = 2.1µm). We were also interested in the distribution of UNC-89 and expected to see a similar pattern for MEC-2. However, when we tagged the endogenous *unc-89* locus at the N-terminus with GFP (fig. S6C), we visualized expression that was largely restricted to the muscles (fig. S6H). With the previous functional results in mind (Fig. 3, A-C and fig. S6, E and G), we conjectured that UNC-89 is expressed in TRNs (table S1, fig. S6B), however, in quantities that cannot be detected above background level if tagged with a single fluorophore. Thus, we sought to visualize the endogenous distribution using multiplexed split FP complementation (*49*) by tagging the endogenous locus with 5x wrmScarlet(11) and coexpressing wrmScarlet(1-10) selectively in TRNs. In this case, we could observe cortical UNC-89 in the cell body of TRNs and a faint expression in the neurites (Fig. 3D), which partially colocalized with MEC-2. This may indicate a possible interaction with MEC-2.

We then directly addressed whether MEC-2 colocalizes with UNC-89 through multivalent interactions of their corresponding PRiM and SH3 domain. Specifically, we tested if the SH3 domain of UNC-89 would co-condensate with MEC-2 along the neurite. To explore this idea, we expressed the UNC-89 SH3 domain fused to GFP together with a MEC-2::mCherry. Strikingly, we observed that both proteins sorted into the same punctae and colocalized along the neurite (Fig. 3E). In contrast, we neither observed colocalization of UNC-89 with the MEC-2(R385H) mutant nor with a GFP fusion to an SH3 domain borrowed from a protein that does not interfere with touch sensation (*spc-1*, see above, Fig. 3, A and E). The picture that is emerging suggests that MEC-2 C-terminus recruits UNC-89 through its SH3 binding domain in TRNs *in vivo*. To further challenge this result, we expressed full-length MEC-2 in body wall muscles, where it is normally not expressed, but contains prominent expression of UNC-89. Consistently, we observed that it formed a characteristic pattern composed of punctae and stripes, that seemingly overlapped and colocalized with the stripes seen for UNC-89 (fig. S6I).

Lastly, we investigated whether MEC-2 and UNC-89 could interact and co-phase separate *in vitro*. We first prepared MEC-2 samples that undergo LLPS and doped them with UNC-89, at molar ratios 1:0.1 and 1:1 (MEC-2:UNC-89). The samples at ratio 1:0.1 showed that UNC-89 can indeed partition into the MEC-2 droplets, both of the WT and the R385H mutant (Fig. 3F). Interestingly, the samples at 1:1 ratio showed complete dissolution of MEC-2 droplets (fig. S7A), indicative of a direct interaction between MEC-2 and UNC-89, that competes against the homotypic interactions driving phase separation. To confirm these results, we co-assembled purified MEC-2 C-terminus with the UNC-89 SH3 domain and tested the interaction by NMR. When we added the unlabeled SH3 to ^15^N labeled MEC-2, we observed a weak but consistent intensity reduction in residues adjacent to the PRiM. This signal intensity reduction was nearly absent in the MEC-2(R385H) mutant, indicating that the mutation abrogates binding (Fig. 3G and fig. S7, B and C). We also performed the reciprocal experiment (unlabeled MEC-2 with a ^15^N labeled UNC-89 SH3 domain) (fig. S7, D and E) and identified that binding occurs between the RT and nSrc loops of the UNC-89 SH3 domain, as expected for a canonical SH3 binding mode (*34*) (Fig. 3H). Taken together, these results show that MEC-2 binds with low affinity to UNC-89 *in vitro* and can contribute to organizing UNC-89 at mechano-electrical transduction sites *in vivo*. The low affinity of SH3 domain interactions could play a functional role, allowing protein-protein interactions to be rapidly remodeled in response to cellular stimuli (*34, 50*).

Finally, we assessed whether UNC-89 partitioning and binding to MEC-2 had an influence in the maturation propensity of MEC-2 droplets *in vitro*. We incubated MEC-2 (WT and R385H) phase separated samples without or with UNC-89 (1:0.1) over 24h and assessed the morphology of the liquid droplets. Strikingly, the MEC-2 WT with UNC-89 sample clearly underwent a liquid-to-solid transition giving rise to the formation of fibrilar-like structures, which was not observed in the case of MEC-2 R285H mutant or in the samples without UNC-89 (Fig. 3I). This result indicates that UNC-89 partitioning and binding to the PRiM in MEC-2 weakens a heterotypic interaction between MEC-2 molecules that kinetically stabilizes the MEC-2 condensates against maturation. This process, known as heterotypic buffering (*19*), is one of the mechanisms that preserves the liquid character of condensates *in vivo*. Although heterotypic buffering is considered key for preventing the liquid to solid transitions thought to be associated with neurodegeneration, we speculate that it here acts as a mechanism to keep MEC-2 primed for undergoing fast functional maturation upon stimulation.

### MEC-2 transmits force during body wall touch

Until here we showed that mature MEC-2 condensates stiffen *in vivo* and endow mechanosensitivity to external touch at distinct mecahnoreceptor sites along the neurite (Fig. 3). We next sought to establish whether or not MEC-2 is able to sustain mechanical load during touch and act directly in transmitting force to the MeT channel. We thus engineered a genetically encoded FRET-tension sensor module (TSMod, (*51, 52*)) into full-length MEC-2 between the stomatin domain and the PRiM (Fig. 4A), with the aim to visualize changes in FRET during touch. Importantly, this insertion did not disrupt localization of MEC-2 (Fig. 4B, fig. S5 and fig. S8A) and preserved partial touch sensitivity (Fig. 4C). We then immobilized these transgenic animals into the body wall chip (*35*) and applied increasing pressure to the side of the animal and released the pressure in one step (Fig. 4D), while imaging FRET signal in a confocal microscope (*53*). In animals bearing the tension sensitive MEC-2 FRET module, we observed a steady decline in the FRET efficiency that is negatively correlated to the pressure applied. Upon sudden pressure release, the FRET index increased again to the same value as before the indentation, indicating that the MEC-2 can reversibly transmit force between the PHB and the C-terminal domain. Consistent with previous reports, tension was highest in regions directly below the actuator and did not propagate into the distal regions of the axon (*54*). In the C-terminal TSMod fusion intended to serve as a pressure-insensitive control (fig. S8B) and in the transgenic animals bearing the MEC-2(R385H) mutation (Fig. 4) we did not observe changes in FRET index with the applied pressure. Importantly, artificially separating the FRET cassette with a 200 amino acid spacer domain led to constitutively low FRET values (fig. S8A), indicating that our FRET measurements report reliable values. Lastly, we asked whether or not the naive, sol-like fraction of MEC-2 would be able to transmit forces. Due to the difficulty of performing the FRET measurements on moving spots under the application of an external pressure, we resorted to the conditionally, C-terminally truncated construct after TEV cleavage. This truncated protein was not able to mature into distinct gel-like condensates and is characterized by high FRET values (fig. S8A), similar to the C-terminal no-force control (fig. S8B). Together, this suggests that MEC-2 is under mechanical tension during touch and therefore is an integral component of the force transmission pathway in *C. elegans* TRNs.

**Figure 4.**
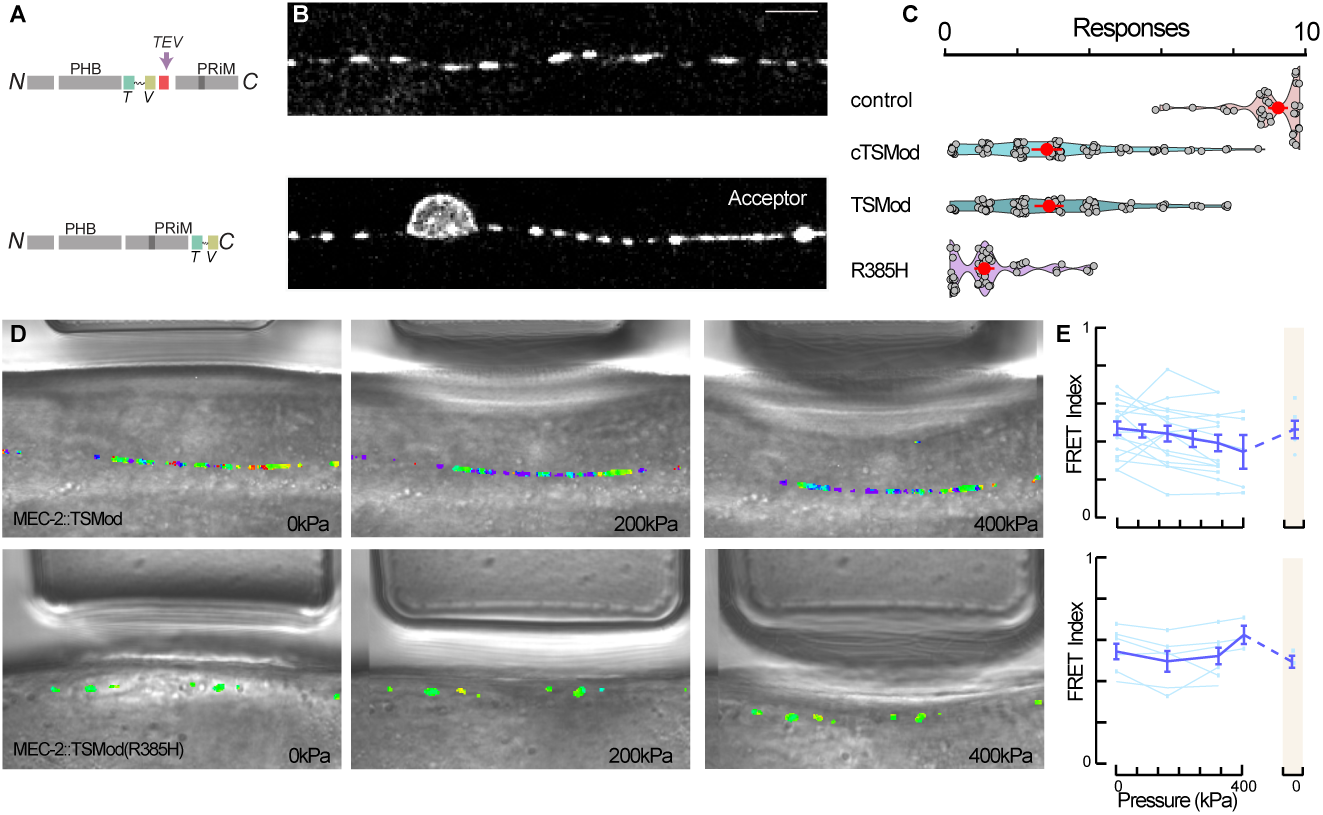
MEC-2 sustains tension during body wall touch. (**A**) Scheme of the FRET-based tension sensing module (TSMod) integrated between the PRiM and the PHB domains (top), or control with free TSMod at the C-terminus (bottom). PRiM, proline-rich motif; PHB, prohibitin domain; TEV, Tobacco etch virus cleavage site; T, donor; V, acceptor fluorophore. (**B**) Representative images of TRNs expressing the internal and C-terminal MEC-2:TSMod fusion protein. Scale bar = 5 µm (**C**) Violin plot of the body touch response derived from N2 wildtype control animals, internal (TSMod) embedded between amino acids 370-371, C-terminal TSMod (cTSMod) fusion and internal TSMod in R385H mutant MEC-2. Circle indicates median, vertical bar indicates SD, N≥ 60 animals. *p*-value derived from Tukey HSD test. (**D**) Representative brightfield images of wildtype (top) and R385H mutant MEC-2::TSMod::MEC-2 (bottom) transgenic animals within the body wall chip under increasing force application (0-200-400 kPa, indicated by the actuator deflection) overlayed with their corresponding FRET index representation. Scale bar=10µm. (**E**) Quantification of the FRET index vs. pressure delivered to the body wall for N=6 animals. Shaded column indicates the resting FRET value at 0 pressure after the pressure is relieved, indicating reversibility of the FRET response.

## Discussion

Despite the efforts in framing a unifying principle of mechanosensation (*55, 56*), the diversity of mechanisms describing the transduction of mechanical stress into biological signals is ever expanding (*53, 57, 58*). We have added a new spotlight on stress-responsive biomolecular condensates that coral mechanosensitive ion channels and host components of the cytoskeleton to activate force gated ion channels.

Our notion that MEC-2 forms BMCs is consistent with previous observations that Stomatin forms higher order oligomers (*4*), while the IDR of MEC-2 directly mediates protein-protein interactions (*7, 59*) and maturation from the liquid into gel-like pools. Because the diameter of *C. elegans* neuronal axons is not constant along their length, we propose that the liquid-like property facilitates transport along the neurite with varying caliber, where material needs to be squeezed through constrictions along the way. Moreover, material can easily be exchanged with the target sites where it assembles into mechanoresponsive clusters, as observed in tight junctions of zebrafish and MDCK epithelia (*21, 22*). We speculate that the increase in relaxation time observed *in vitro* (Fig. 2 and fig. S3) has profound consequences on their mechanical function as a tension sensor (*15*). Once at their target, the MEC-2 condensates undergo liquid-solid transition such that they are able to sustain mechanical stress over long timescales during body wall touch, which is providing a focus for force transmission to the ion channel. In addition, the solid transition might itself produce a stress that reshapes the neuronal membrane (*14, 60*). If additional factors and proteins promote the maturation is an open question for future experiments.

The data presented here point towards a mechanism by which MEC-2/Stomatin condensates host UNC-89/Titin/Obscurin with its SH3 domain, however, it is unlikely that UNC-89 is directly involved in force transfer. Rather, we propose that the SH3 binding to MEC-2 accelerates a rigidity percolation through crosslinking of the condensate with homotypic and heterotypic interactions that lead to a dynamic arrest and changes in material properties. This increase in the timescale of crosslink failure determines the efficiency of force transmission during touch. Indeed, a change in a single residue within the MEC-2 PRiM that determines the binding to UNC-89 changes the percolation threshold with consequences on material properties, neuronal activity and the touch response. Our results thus demonstrate for the first time a physiological role of a rigidity percolation in mechanotransduction within a biomolecular condensate.

Our results show that UNC-89/Titin/Obscurin is expressed and functional in neurons (Fig. 3). We observed expression in touch receptor neurons, but also motor neurons and amphid cells of *C. elegans*, confirming recent neuronal RNAseq data (*41*). The mammalian homolog ObscurinB is also expressed in the brain (*47*), and might have implications for neurodegenerative diseases. Several reports indicate a potential role of Titin in motor neuron diseases. Copy number variations and single nucleotide polymorphism with potential pathogenicity have been identified in patients with spontaneous amyotrophic lateral sclerosis, a fatal motor neuron disease (*61, 62*). It was hypothesized that these changes in motor neuron function were induced by aberrant Titin expression in muscles (*62*). Our data showing functional Titin expression in neurons, including motor neurons of *C. elegans*, motivates to revisit the neuronal expression pattern in mammals and the contribution of Titin in motor neuron diseases in humans.

Taken together, we identified a mechanism by which the coacervation of a scarce SH3 domain induces a rigidity percolation in a disordered network, that confers mechanical stability and mechanotransmission during external touch. Future work needs to address how mechanical force is transmitted to the percolated networks and modulates the UNC-89/Titin/Obscurin interaction within the condensate.

## Acknowledgments

We thank the NMSB and SLN labs for discussions and suggestion throughout the work and for use of their microscopes. We thank the ICFO BIL and NFL for support with animal maintenance and SU8 lithography, respectively. We thank Li-Chun Lin and Shadi Karimi for help with MATLAB code and fabrication of microfluidic devices, respectively. We thank the ICTS NMR facility, managed by the scientific and technological centers of the University of Barcelona (CCiTUB), for their help in NMR. We thank the IRB Barcelona advanced digital microscopy facility, for their help with microscopy experiments. We thank Martin Chalfie, Miriam Goodman, Martin Harterink, Carlo Carolis and the CGC (National Institutes of Health - Office of Research Infrastructure Programs (P40 OD010440)) for providing reagents; Ben Lehner and Julian Cerón for sharing their RNAi libraries.

## Authors contribution

NSC: animal husbandry, molecular biology, CRISPR, CRE recombination, optogenetic and behavioral experiments, FRET and FRAP assays, calcium imaging, data analysis and manuscript writing. BM: NMR and LLPS characterization. CGC and MR: LLPS characterization and FRAP. FCC: optical tweezer microrheology. IR: particle tracking and MSD calculations. MP: molecular biology. XS: study conceptualization. MK: study conceptualization, acquisition of funding, data analysis and manuscript writing.

## Data and materials availability

All data is available in the manuscript or the supplementary materials.

## Supplementary material

Other Supplementary Materials for this manuscript include the following

### Materials and Methods

#### C. elegans culture

Animals were maintained on Nematode Growth Medium (NGM) plates seeded with *Escherichia coli* OP50 bacteria. Age-synchronized young adult animals were used for all the experiments and handled as described (*64*). All strains generated in this study are listed in table S2.

#### Molecular biology and transgenesis

All plasmids listed in table S3 were generated using the Gibson assembly method. All coding sequences were verified by sequencing.

#### Expression of WT and R385H *mec-2* and *unc-89* for *in vitro* purification

Wild-type complementary DNA ranging from position 371 to 481 of MEC-2, which includes the C-terminal domain, was subcloned into a pCoofy expression vector (donated by Carlo Carolis lab) containing an N-terminal polyhistidine affinity tag and a NusA solubility tag for posterior MEC-2 purification, giving to pNS66. This plasmid was used as a template to incorporate the R385H single point mutation (*u26* allele) in MEC-2 by site-directed mutagenesis, giving to pNS73. Wild-type complementary DNA encoding 61-128 amino acids of UNC-89, which includes the SH3 domain, was subcloned into a pET-24 expression vector (ordered from Twist Bioscience) with an N-terminal polyhistidine affinity tag for posterior UNC-89 purification, giving to pNS75. A previous version including 1-454 amino acids of UNC-89 led to unfolded purified protein.

#### Expression of WT and R385H *mec-2*::mCherry

Wild-type complementary DNA encoding full-length MEC-2 (A isoform) was subcloned into pBCN27 (*65*) to replace puromycin resistance gene, and fused to mCherry, generating pMK8. Site-directed mutagenesis was used to introduce the R385H single point mutation (*u26* allele) in MEC-2, giving pMK9. Both plasmids were integrated by the MosSCI method (*66*) in the EG6699 strain, that contains compatible MosSCI landing sites in Chr. II, leading to MSB87 and MSB88, respectively.

#### Generation of the *u37* allele in *mec-2*

The *mec-2(u37)* allele (W119Stop), which introduces a premature stop codon, was reproduced by CRISPR/Cas9 genome editing as described in (*67*). Two crRNAs were designed to cut few base pairs before the target site together with a donor consisting of a ssODN with 35 basepair (bp) homology arms flanking the polyspacer adjacent motif (PAM) sequence, the desired single point mutation and 5 other silent mutations to facilitate the posterior screening of the edit by PCR. Briefly, the Cas9-crRNA-tracrRNA RNP complex and the homology repair template (HDR) were assembled in Mili-Q water, together with the Cas9 complexes and HDR for the marker gene *dpy-10*, to introduce the semidominant *cn64* allele. 20-30 young adult hermaphrodites were injected with the CRISPR mix and recovered onto individual plates. After 3 days cultured at 25°C, the progeny was screened based on the dpy or roller phenotype and singled onto individual plates. Mothers were lysed, screened by PCR for the corresponding edit and verified by sequencing. Sequences of crRNAs and ssODN donors are provided in table S4.

#### Expression of *mec-2* in muscles and hypodermis

*mec-2* full-length cDNA and mCherry fluorescent protein were amplified from pMK8 and pNS10, respectively, and cloned as translational fusion under the *myo-3* muscle promotor from pNS60. The resulting plasmid, pNS68, was injected in MSB523 animals at 10 ng/ul, as well as the control plasmid pMK23 (*myo-3*p::mCherry), leading to MSB938 and MSB937, respectively. For *mec-2* expression in hypodermis, the *wrt-2* hypodermal promotor was amplified from genomic DNA (1376 bp), *mec-2*::mCherry::coLOVpep was amplified from pNS13 and cloned into pNMSB35 backbone giving pNS70. It was injected at 30 ng/ul leading to MSB991.

#### Generation of the FRET constructs

The TSMod cassette containing mTFP, 40-amino acid-long flexible linker, mVenus and a TEV protease site, was amplified from pMG319 (*68*) and inserted between 370-371 amino acids of *mec-2*, leading to pNS2. This plasmid was used as a template to introduce the R385H mutation by site-directed mutagenesis to yield pNS24. The TSMod cassette was also inserted at the C-term of *mec-2* in the plasmid pMK13. These plasmids were integrated by the MosSCI method (*66*) in Chr II in strains with *mec-2(u37)* background in the endogenous copy (Chr X), generating MSB341, MSB357 and MSB74, respectively. The low FRET construct was made by replacing the TSMod cassette in pNS2 with a mTFP-TRAF-mVenus cassette derived from pMG352 (*68*), which constitutively separates the donor and acceptor fluorophores, and it was injected as extrachromosomal array giving to MSB907. A TEV protease site was fused to mCherry through a spliced leader SL2 (gpd-2-gpd-3) under TRN-specific *mec-17* promotor, giving to pMK97. It was injected into MSB341 leading to MSB403.

#### Promotor trapping of *unc-89*

The *unc-89* promotor expression vector, was generated by amplifying 4 kb upstream to *unc-89* gene and the first three exons, including the SH3 domain, from *C. elegans* genomic DNA (table S3). It was transcriptionally fused to GFP through a spliced leader SL2 (gpd-2-gpd-3), generating pNS49, which was injected in MSB87 animals, leading to MSB656.

#### Generation of (mEGFP(loxP)::AID knock-in)*unc-89*

The Nested CRISPR/Cas9 genome editing (*69*) was used to knock-in mEGFP at the *unc-89* gene. Two crRNAs were used to cut the N-term of *unc-89* and it was repaired by 200 bp ssODN containing parts 1 and 3 of mEGFP including a loxP within a synthetic intron of the mEGFP, along with a flexible linker and a degron site (AID). A new PAM site and a protospacer sequence was inserted in the first fragment to allow the in-frame insertion of the remaining sequence mEGFP2, designed as an IDT gBlock. For the second step, the same universal crRNA mentioned in (*69*) was used to make the double stranded break (see table S4). The correct in-frame insertion of the full length mEGFP was sequence verified and correct UNC-89 expression was checked by green fluorescence expression in muscles. The knock-in was done on top of MSB87, generating MSB523.

#### Generation of the conditional and constitutive *unc-89* knock-out

*unc-89* knock-out was generated by CRISPR/Cas9 genome editing using two crRNAs to cut exon 3 of *unc-89* and a 108 bp ssODN that led to a frame-shift and absence of the largest isoforms (a,b,e,f,k,l,m,n,o) of *unc-89*, see table S4. It was done on MSB523 mEGFP(loxP)::degron::*unc-89*, generating MSB590. Animals were verified by sequencing and by absence of green fluorescence in muscles. The resulting animals move normally but show a slight delay in development and body size (fig. S6D).

#### CRE/loxP and AID degradation

Tissue specific *unc-89* knock-out was generated by inserting an in-frame loxP site at the C-term of the *unc-89* largest isoforms by CRISPR/Cas9 genome editing. Two crRNAs were used to cut the C-term of *unc-89* and it was repaired by a 129 bp ssODN that carried a loxP and 5 silent mutations for posterior screening by PCR (see table S4). It was injected in MSB523 (mEGFP(loxP)::AID::*unc-89*) giving to MSB930. The correct in-frame insertion was sequence verified and correct UNC-89 expression was checked by green fluorescence expression in muscles. MSB930 was crossed to MSB926, which carries the TRN-specific *mec-17*p::CRE and the *mec-4*p::TIR (*53*) leading to MSB953. Alternatively, MSB930 was crossed to MSB933 which carries the panneuronal (*rgef-1*)p::CRE leading to MSB941.

#### Tagging of *unc-89* SH3 domain

The plasmid to tag *unc-89* SH3 domain (63-127 amino acids) was generated by amplifiying the SH3 motif from N2 genomic DNA and cloning it under the TRN-specific *mec-18* promotor from pMK105. It was fused to GFP with a 5 amino acids linker from IR83 giving to pNS41. For *spc-1* SH3 tagging, the *mec-17* promotor and *spc-1* SH3 motif were taken from the pMK32 backbone and fused to GFP with a 3 amino acids linker from pDD282 giving to pMK101. They were injected at 20 ng/ul giving to MSB493 and MSB544, respectively.

#### Prediction of phase separation behavior

The primary sequence of the MEC-2 A isoform was imported into AlphaFold2 and transform restraint Rosetta for structure prediction using the standard parameters for folding. Prion like sequences were predicted using the methodology described in reference (*27*) with the *C. elegans* proteome as a background sequence. Phase separation index was calculated using software presented in (*28*).

### *In vitro* assays

#### Protein Expression and purification

To obtain unlabelled MEC-2 protein, *E. coli* B834 (DE3) cells were transformed with the MEC-2 C-terminal (371-481) plasmid (pNS66). The cells were grown in LB medium at 37 °C until OD=0.6 and induced by the addition of isopropyl *β*-D-1-thiogalactopyranoside (IPTG) to a final concentration of 1 mM. The cultures were grown overnight at 25 °C. After 30 min centrifugation at 4000 rpm, the cells were resuspended in 50 mM Tris, 50 mM NaCl and 1 mM dithiothreitol (DTT) buffer at pH=7.4. The cells were lysed by sonication and centrifuged for 30 min at 20000 rpm. The supernatant was loaded in a nickel affinity column (HisTrap HP 5mL, Cytiva) and eluted with a gradient from 0 to 500 mM imidazole.The histidine affinity tag was cleaved with 3C protease through dialysis for 2 h in cleavage buffer (50 mM Tris-HCl, 50 mM NaCl, 1 mM DTT, pH 7.4). 8 M urea was added in order to separate NusA and MEC-2. The reverse nickel column was run with 8 M urea to remove the cleaved tag and uncleaved protein. After loading, MEC-2 was eluted with a buffer containing 50 mM Tris-HCl, 300 mM NaCl, 20 mM imidazole, 1 mM DTT and 8 M urea at pH=8.0. The eluted protein was injected in a size exclusion Superdex 75 (Cytiva), running in 20 mM sodium phosphate buffer with 1 mM TCEP and 0.05% NaN3 at pH 7.4. The fractions with protein were joined and concentrated to 400 µM, fast frozen in liquid nitrogen and stored at -80 °C.

To obtain unlabelled UNC-89 protein, *E. coli* B834 (DE3) cells were transformed with the UNC-89 SH3 (59-128) plasmid (pNS75). The cells were grown in LB medium at 37 °C until OD=0.6 and induced by the addition of IPTG to a final concentration of 1 mM. After growing 4 h at 37 °C, the cells were centrifuged for 30 min at 4000 rpm and resuspended in 50 mM Tris and 50 mM NaCl buffer at pH=7.4. The cells were lysed by sonication and centrifuged for 30 min at 20000 rpm. The pellet was washed twice with washing buffer (PBS, 1 mM DTT, 500 mM NaCl, 1% TritonX-100, PIC, PMSF, DNAse and RNAse, at pH 7.4). The pellet was resuspended in resuspension buffer (25 mM Tris-HCl, 8 M urea, 500 mM NaCl, 10 mM imidazole, pH 8.0) and centrifuged for 30 min at 20000 rpm. The supernatant was loaded in a nickel affinity column (HisTrap HP 5 mL, Cytiva) and eluted with a gradient from 0 to 500 mM imidazole. The histidine affinity tag was cleaved with 3C protease through dialysis for 2 h in cleavage buffer (50 mM Tris-HCl, 200 mM NaCl, 1 mM DTT, pH 8). The reverse nickel column was run to remove the cleaved tag and uncleaved protein. After loading, UNC-89 was eluted with a buffer containing 50 mM Tris-HCl, 300 mM NaCl, 20 mM imidazole, 1 mM DTT and 8 M urea at pH=8.0. The eluted protein was injected in a size exclusion Superdex 75 (Cytiva), running in 20 mM sodium phosphate buffer with 1 mM TCEP and 0.05% NaN_3_ at pH 7.4. The fractions with protein were joined and concentrated to 1 mM, fast frozen in liquid nitrogen and stored at -80 °C.

Isotopically ^15^N/^13^C- and ^15^N-labelled proteins were produced by growing transformed *E. coli* B834 cells in M9 minimal medium containing 1 g·L^-1^ of ^15^N-NH_4_Cl and 2 g·L^-1^ of ^13^C_6_-D-glucose.

#### NMR experiments

##### Backbone assignment

NMR experiments were recorded at 278 K on a Bruker Avance NEO 800 MHz spectrometer or a Bruker Avance III 600 MHz, both equipped with a TCI cryoprobe. A 400 µM ^15^N, ^13^C double labelled MEC-2 (371-481) sample in NMR buffer (20 mM sodium phosphate (pH 7.4), 1 mM TCEP, 0.05 % (w:w) NaN_3_) was used for backbone resonance assignment. A series of 3D triple resonance experiments were recorded, including the BEST-TROSY version of HNCO, HN(CA)CO, HNCA, HNCACB, and HN(CO)CACB (*70*). Chemical shifts were deposited in BMRB (ID:51491). Secondary structure propensities were derived from the H, N, C’, C*^α^* and C*^β^* chemical shifts measured by using solution state NMR and using the δ2D software (*29*). A 1200 µM ^15^N, ^13^C double labelled UNC-89 SH3 domain (59-128) sample in NMR buffer (20 mM sodium phosphate (pH 7.4), 1 mM TCEP, 0.05 % (w:w) NaN_3_) was assigned using the same NMR experiments as described above. Chemical shifts were deposited in BMRB (ID:51490).

##### Binding mapping

Chemical shift perturbations (CSP) and signal intensity changes (I/I_0_) were extracted by measuring ^1^H-^15^N correlation spectra of ^15^N, ^13^C double labelled MEC-2 (371-481) with 10 molar equivalents of UNC-89 SH3 domain, and vice versa. Data analysis was performed with CcpNmr V3 (*71*).

#### Sample preparation for *in vitro* experiments

All samples were prepared as follows. First, a buffer stock solution consisting of 20 mM sodium phosphate, 1 mM TCEP and 0.05 % NaN_3_ was pH adjusted to 7.4 and filtered using 0.22 µm sterile filters (Buffer Stock). A 5 M NaCl solution in the same buffer was also pH adjusted to 7.4 and filtered (Salt Stock). Then, the protein samples were thawed from -80 °C on ice, pH adjusted to 7.4 and centrifuged for 5 minutes at 15000 rpm. The supernatant (Protein Stock) was transferred to a new Eppendorf tube and the protein concentrations were determined by their absorbance at 280 nm. The indicated samples were prepared by mixing the right amounts of Buffer Stock, Protein Stock and Salt Stock to reach the desired final protein and NaCl concentrations.

#### Apparent absorbance as a function of temperature

Absorbance of the samples was measured at 350 nm (A_350_*_nm_*) using 1 cm pathlength cuvettes and a Cary100 ultraviolet–visible spectrophotometer equipped with a multicell thermoelectric temperature controller. The temperature was increased progressively from 10 to 60 °C at a ramp rate of 1 °C/min. The cloud temperatures (Tc) were determined as the maximum of the first order derivatives of the curves.

#### Differential interference contrast microscopy

1.5 µL of sample was deposited in a sealed chamber comprising a slide and a coverslip sandwiching double sided tape (3 M 300 LSE high-temperature double-sided tape of 0.17 mm thickness). The used coverslips were previously coated with PEG-silane following the published protocol in ref. (*72*). The DIC images were taken using an automated inverted Olympus IX81 microscope with a 60x/1.20 water UPlan SAPo objective using the Xcellence rt 1.2 software.

#### *In vitro* confocal fluorescence microscopy

MEC2 WT or R385H were labeled with Alexa Fluor 647 and UNC-89cysmutant with DyLight 488, following provider’s instructions (Thermo Fisher Scientific). The samples for fluorescence microscopy were prepared as previously described but containing 1 µM of labeled protein molecules. UNC-89cysmutant plasmid was generated by directed mutagenesis on top of pNS75 backbone to incoporate a Ser to Cys change in amino acid 62 for posterior fluorescence labeling giving to pNS77.

Fluorescence microscopy images and FRAP experiments were recorded using a Zeiss LSM780 confocal microscope system with a Plan ApoChromat 63x 1.4 oil objective. For the FRAP experiments, 10 or 11 droplets of similar size were selected for MEC2-R385H or MEC2-WT, respectively. The bleached region was 30% of their diameter, and the intensity values were monitored for ROI1 (bleached area), ROI2 (entire droplet) and ROI3 (background signal). The data was fitted using the EasyFrap software (*73*) to extract the kinetic parameters such as the half-time of recovery and the mobile fraction.

#### Optical tweezer mechanics measurements

Droplet assembly was initiated by mixing Salt Stock Buffer for a final salt concentration of 2 M and a final protein concentration of 370 µM, adjusted to a final volume of 10 µL with Stock Buffer. The Salt Stock Buffer contained 1 µm polystyrene microbeads (Micromod 01-54-103, PEG300 modified) diluted 1:1000 (9.3 · 10^7^mL*^−^*^1^). Bottom glass dishes (GWST-5040, WillCo Wells) were coated with PDMS as described elsewhere (*74*). After curing for 1 h at 65°C, a 5x5 mm hole was drawn out and the glass surface was treated with PLL(20)-g[3.5]-PEG(2) (30 min, 0.5 mg/mL, SuSoS) to yield protein condensation with a spherical shape (Fig. 2E). A 10 µl drop of the aforementioned solution was added in the middle of the cavity, which was later closed with a 1x1 inch coverglass very gently to avoid air bubbles. An optical trap was created at the focal plane of a water-immersion objective (60x, NA=1.2, Plan Apo, Nikon), using an optical micromanipulation unit (Sensocell, Impetux Optics) coupled to the rear epi-fluorescence port of an inverted microscope (Nikon Eclipse Ti2). Optical traps were equipped with a light momentum force detection module (Sensocell, Impetux Optics) that substituted the microscope brightfield illumination condenser. The trap stiffness, k*_OT_* (pN), was obtained by performing a fast scan over the trapped particle and fitting a line over the linear range, i.e. -100 nm < x < 100 nm, as described elsewhere (*75*). After that, the bead was brought into contact with the protein droplet surface (Fig. 2I). Upon contact, the optical trap measured a steep increase in the force experienced by the bead against the droplet surface. The optical trap was programed to perform different trajectories using the software provided by the manufacturer (LightAce, Impetux Optics). For the Lissajous curves displayed in Fig. 2J, sinusoidal pushing onto the droplet was performed at frequencies f=0.2,1,4,10 Hz, with an amplitude of A=±100 nm, while the actual bead position was obtained as x*_bead_* = x*_trap_*-F/k*_OT_* . Stress relaxation measurements (fig. S3B) were obtained by applying a square oscillation (0.2 Hz, A=±100 nm, Fig. S3B). The force relaxation curves were fitted with an exponential decay to determine the time constant, τ (s), and the droplet stiffness, k = f*_p_*/δ (µN/m) (fig. S3, C and E, and fig. S4, O and P). Mechanical measurements on protein droplets were repeated at t=4, 24 and 48 h. Microchambers containing the droplets and the polystyrene microspheres were kept at room temperature meanwhile. Data processing was carried out with custom scripts in Matlab.

### Behavioral assays

#### Gentle body touch assays

Gentle body touch assays were carried out as described elsewhere (*76*). An eyebrow hair was used to gently touch 20-30 young adult worms for ten times with alternative anterior and posterior touches (five each). Unless otherwise stated, these experiments were repeated at least 3 different days to determine an average response and SD. The results of the touch assays are included in table S5.

#### TRN-specific RNAi feeding

RNAi bacterial clones were obtained from Ahringer (*42*) (*mec-2*, *mec-4*, *ced-5*, *sem-5*, *toca-1*, *tbc-18*, *sdpn-1*, *F49E2.2*, *unc-73*, *spc-1*, *mlk-1*, *abl-1*, *sma-1*, *plc-1*, *itsn-1*, *nck-1*, *magu-3*, *ccb-1*, *C46H3.2*, *prx-13*, *erp-1*, *ephx-1*) or Vidal (ORFeome-Based) (*77*) (*B0303.7*, *abi-1*, *magu-2*, *unc-89*, *stam-1*, *lin-2*, *F42H10.3*, *amph-1*, *dbn-1*, *C36E8.4*, *T04C9.1*, *unc-57*, *Y106G6H.14*, *ced-2*) libraries, donated by Cerón and Lehner labs. First, clones were verified by colony PCR and sequencing. NGM plates were supplemented with 6 mM IPTG (I1001-25 Zymo Research) and 50 µg/mL ampicillin. Unseeded feeding plates were completely dried in a laminar airflow hood and kept in the dark at 4°C. RNAi bacterial clones were grown in 4 mL LB with 50 µg/mL ampicillin. Next day, each plate was inoculated with 200 µL of the corresponding bacterial culture and dried for 1 h under the hood. The expression of dsRNA was induced in the presence of IPTG overnight at room temperature, or in an incubator at 37°C for 4 h, in the dark. Then, 6 gravid hermaphrodites (TU3403 (*44*)) were transferred onto the plates and grown at 25°C for 48 h. The progeny were tested for body touch sensitivity at young adult stage, a total of 40 worms in 2 different days. Importantly, wildtype animals are insensitive for neuronal RNAi due to the lack of dsRNA transporter in neurons. Thus, animals were sensitized to RNAi through a TRN-specific SID-1 rescue construct (*mec-18*p::*sid-1*(+); (*44*)) in a systemic RNAi mutant *sid-1*(qt2) background.

#### Auxin-induced degradation experiment

Auxin plates were prepared as described elsewhere (*78*), by adding 250 mM stock of 1-naphthaleneacetic acid (NAA Auxin, Sigma Aldrich 317918) dissolved in 95% ethanol to cooled NGM before pouring the plates at a final concentration of 1 mM auxin. Plates were dried out under a laminar airflow hood, seeded with 10X concentrated *E. coli* OP50 and dried for 1 h under the hood. Next day, 6 gravid hermaphrodites were transferred onto the plates and grown at 25°C for 48 h. The progeny were tested for body touch sensitivity at young adult stage.

Calcium imaging of TRNs from microfluidically-immobilized animals after body wall touch

#### Device preparation

Devices were replica-molded from SU8 photolithography mold as described previously (*35*).

#### Animal loading into a microfluidic trap

Loading of the animals in the body wall chip was performed as described in detail elsewhere (*79*). Briefly, 2-3 young adult animals were transferred to a M9-filtered droplet. Then, worms were aspirated through a SC23/8 gauge metal tube (Phymep) connected to a 3 mL syringe (Henke Sass Wolf) with a PE tube (McMaster-Carr). Once the tube was inserted in the inlet of the chip, the animals were loaded on the waiting chamber by applying gentle pressure with the syringe. In general, animals were oriented head-first.

#### Calcium imaging

*In vivo* calcium imaging of TRNs was performed by positioning the worm-loaded microfluidic device in a Leica DMi8 microscope with a 40x/1.1 water immersion lens, Lumencor Spectra X LED light source, fluorescence cube with beam splitter (Semrock Quadband FF409/493/573/652) and a Hamamatsu Orca Flash 4 V3 sCMOS camera. Cyan-488 nm (≈6.9 mW) and yellow-575 nm (≈12.6 mW) illuminations were used to excite the green and red fluorescence of the TRN::GCaMP6s and mtagRFP, which was used to correct possible artifacts from animal movement and TRN identification. The incident power of the excitation light was measured with a Thorlabs microscope slide power meter head (S170C) attached to PM101A power meter console. Emission was split with a Hamamatsu Gemini W-View with a 538 nm edge dichroic (Semrock, FF528-FDi1-25-36) and collected through two single band emission filters, 512/525 nm for GCaMP (Semrock, FF01-512/23-25) and 620/60 for mtagRFP (Chroma, ET620/60m). The emission spectra was split by the image-spliter, allowing different exposure times for each signal. For mechanical load application to body wall of the animal, the stimulation channel was connected to a piezo-driven pressure controller (OB1-MK3, Elveflow) as described (*80*). To follow calcium transients, videos were taken at 10 frames-per-second with 80 ms exposure time, using the master pulse from the camera. The camera SMA trigger out was used to synchronize the stimulation protocol in Elveflow sequencer, which consisted on 20 s pre-stimulation, 2 s stimulation (2500 mbar buzz) and 40 s post-stimulation.

#### Calcium analysis

Images were processed using MATLAB in-house procedures to extract GCaMP signal intensity (*53*). First, the TRN was manually labelled based on the mtagRFP calcium insensitive channel. The position was automatically tracked in the following frames and used to extract the GCaMP intensity. A smooth filter (moving average filter) was applied. The calcium sensitive signal was normalized to the first 100 frames pre-stimulation (F-F_0_/F_0_) and the results were plotted in Python.

### Fluorescence resonance energy transfer (FRET)

#### Data acquisition

FRET imaging of worms loaded within the body wall chip as described above was performed on a Leica DMI6000 SP5 confocal microscope using the 63x/1.4 NA oil immersion lens. As described in detail in (*68*), three images were collected: the direct donor (mTFP2) excitation and emission, donor excitation and acceptor emission and direct acceptor (mVenus) excitation and emission. mTFP2 was excited with 458 nm (≈9 µW), mVenus with 514 nm (≈4 µW) line of an Argon ion laser at 80% and 11% transmission respectively (25% power). The incident power of the excitation light was measured with a Thorlabs microscope slide power meter head (S170C) attached to PM101A power meter console. A single set of images was collected before and after pressure delivery, while recording images for each pressure. Due to defocussing immediately after the pressure delivery, manual refocussing was necessary to keep the focus in the plane of the MEC-2 clusters.

#### Data analysis

Due to the spectral overlap of the donor and the acceptor, the resulting FRET images are contaminated with donor bleedthrough and acceptor cross-excitation. To eliminate this spurious signal, a linear unmixing procedure as a bleedthrough correction was employed. The images were first bleedthrough corrected with a factor that was predetermined prior to each experiment with animals that express either fluorophore alone (for details about the procedure, see Ref. (*53, 68*)). The corrected FRET channel was then normalized by the sum of the background corrected donor channel and the corrected FRET channel on a pixel-by pixel basis. With the aim to eliminate pixels outside of the region of interest and ubiquitous autofluorescence inherent to living *C. elegans*, we applied a mask on the acceptor channel to separate the MEC-2 clusters from the background.

### Fluorescence recovery after photobleaching (FRAP)

#### Data acquisition

FRAP imaging was performed on a Leica DMI6000 SP5 confocal microscope using the 63x/1.4 NA oil immersion lens. Animals were imaged live in 3 mM levamisole on 5-6% agarose pads. The FRAP protocol consisted on 5 frames pre-bleach (every 371 ms), 5 frames of bleach (every 344 ms), 10 frames post-bleach 1 (every 371 ms) and 10 frames post-bleach 2 (every 20 s). mCherry from MSB547 or mVenus from MSB403 were excited with a 594 or 514 nm line of an Argon ion laser, respectively, at 5% of total power, except for the bleach step for which it was set at 100%. For bleaching a small ROI within MEC-2 punctae, same protocol was applied, except for the bleach step where 60% 594 nm line of Argon ion laser was used. For bleaching a small ROI within MEC-2 condensates in hypodermis, the protocol consisted on 5 frames pre-bleach (every 97 ms), 5 frames of bleach (every 66 ms), 10 frames post-bleach 1 (every 1 s) and 10 frames post-bleach 2 (every 20 s).

#### Data analysis

Images were pre-processed using ImageJ/Fiji. First, ROI1 was manually drawn in the bleached area and used to track the intensity measurement in the following frames. The same was done for the total fluorescence area (ROI2) and the background area (ROI3). This data was processed by the easyFRAP online tool (*73*) to compute the normalized recovery curves using full scale normalization, which corrects for differences in the starting intensity in ROI1, differences in total fluorescence during the time course of the experiment and differences in bleaching depth. For the analysis of the conditions where we bleached a small region within MEC-2 punctae or MEC-2 condensate in hypodermis, an extra step of normalization to the bleaching rate (considering first frames of pre-bleach from another ROI in a different punctae/condensate) was added, since the intensity of the whole punctae/condensate decreased during bleaching and could not be used to correct for fluorescence changes during the course of the experiment.

### Confocal microscopy

Fluorescence images were taken using an inverted confocal microscope (Nikon Ti2 Eclipse) with a 60x/1.4 NA oil immersion lens. Animals were imaged live in 3 mM levamisole on 5-6% agarose pads. mCherry was excited using the 561 nm laser, 20-30% power intensity and transmitted through a 594 nm emission filter. Exposure time was 100-200 ms, depending on the strain to image. GFP was excited with a 488 nm laser, 20-40% power intensity and transmitted through a 521 nm emission filter. Exposure time was 100-200 ms.

### Tracking of MEC-2 along TRN

Fluorescently labelled MEC-2 was imaged in the axons of the TRNs using a Leica DMi8 microscope with a 63x/1.4 oil immersion lens. Imaging was performed at a frame rate of 100 ms. The naive pool was imaged in close proximity to the cell body and the mature pool at the perifery of the axon. MEC-2 trajectories of the mobile pool were extracted using MTrackJ (*81*) and the trajectories for the immobile pool were obtained with the ImageJ Plugin TrackMate (*82*). The resulting trajectories were further analysed using a Python script to compute the mean squared displacement defined as

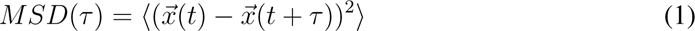

The average was taken over the time and the ensemble of the measured trajectories.

### Interpunctum interval analysis

For calculating the *mec-2* interpunctum interval, a 160-190 µm length ROI was drawn in TRN axon images in ImageJ. A threshold value and background subtraction were applied to remove the particles outside TRNs. The ImageJ particle counting tool was used to infer the position of each particle and the difference between them was derived. The resulting values were used to calculate the mean difference using Python.

### Statistics and reproducibility

No statistic method was used to predetermine sample sizes. Statistical methods, repeatably of experiments and N values are indicated within the figure legends.

---

**Fig. S1.**
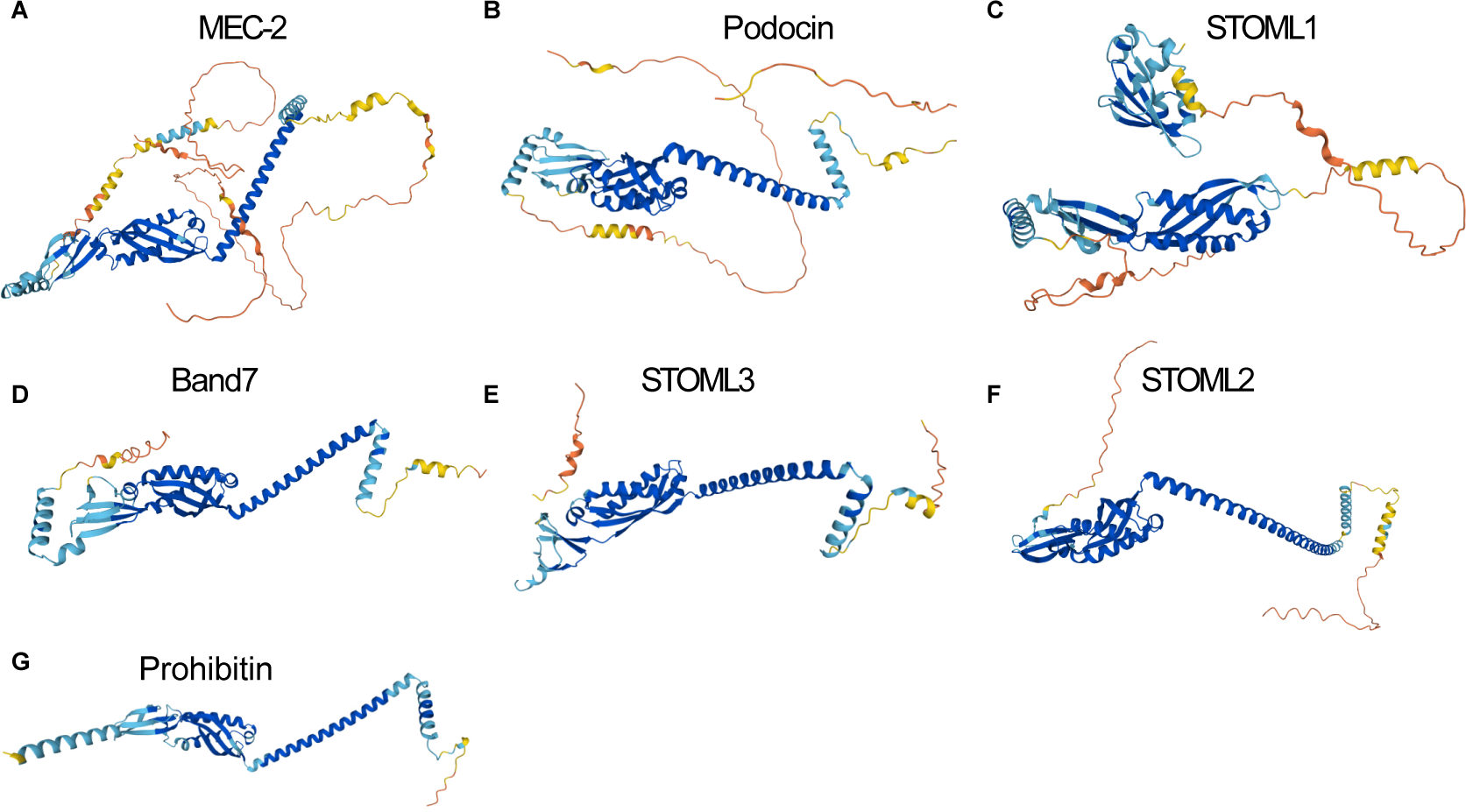
Stomatin proteins structure. Structural prediction of (**A**) worm MEC-2 (Q27433), (**B**) human Podocin (Q9NP85), (**C**) human STOML1 (Q9UBI4), (**D**) Band7 (P27105), (**E**) human STOML3 (Q8TAV4), (**F**) human STOML2 (Q9UJZ1) and (**G**) human Prohibitin (P35232) generated with AlphaFold2.

**Fig. S2.**
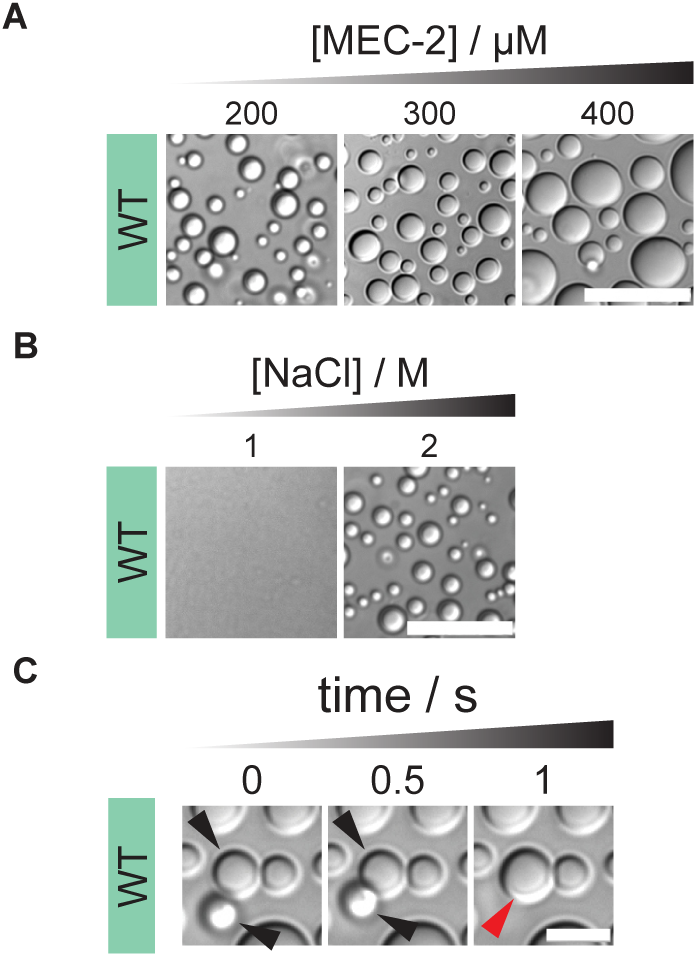
MEC-2 liquid droplets characterization *in vitro*. (**A**) DIC microscopy images of MEC-2 WT at increasing protein concentrations of 200, 300 and 400 µM, with 2 M NaCl at 37 °C. Scale bar = 20 µm. (**B**) DIC microscopy images of 200 µM MEC-2 WT with 1 and 2 M NaCl at 37 °C. Scale bar = 20 µm. (**C**) DIC microscopy images showing fusion events of 300 µM MEC-2 WT with 2 M NaCl at 37°C. Scale bar = 5 µm.

**Fig. S3.**
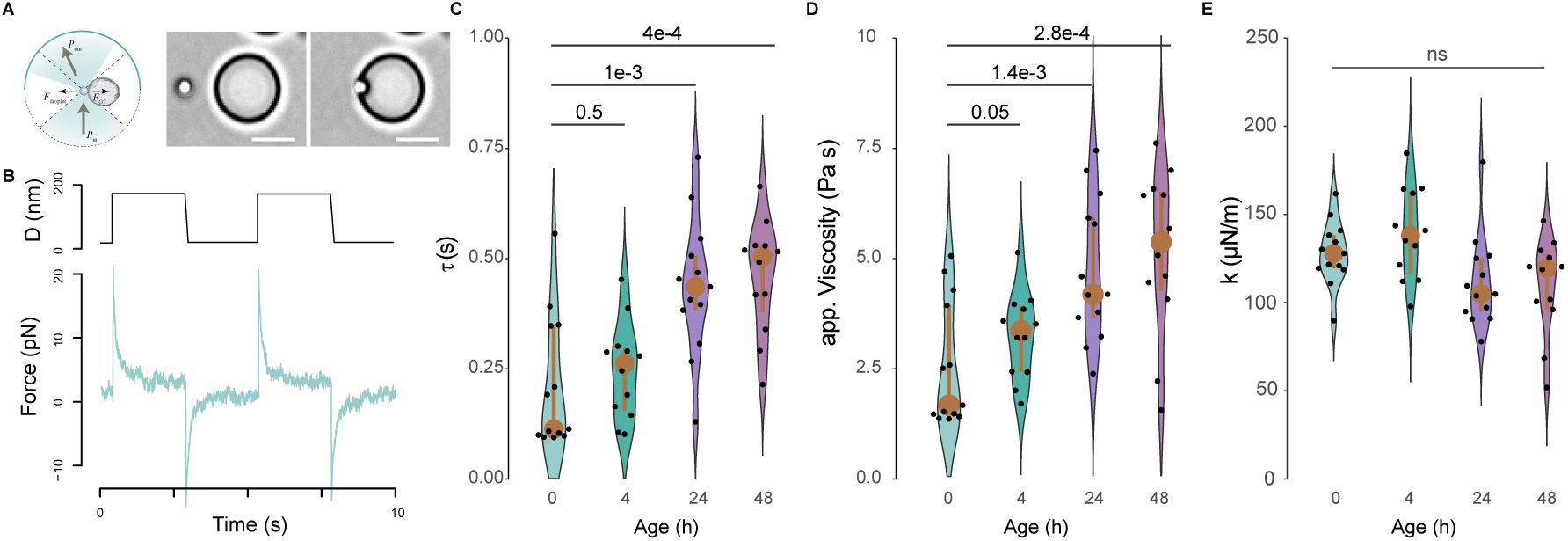
Viscoelastic maturation of MEC-2. (**A**) Scheme of the optical tweezer based indentation assay, during which a trapped microsphere is driven onto an immobilized droplet. Two representative pictures showing the sphere before and after droplet contact. Scale bar = 5 µm. (**B**) Representative force-time signal of a typical indentation test. Upper graph indicates trap trajectory, lower graph stress relaxation. (**C**) Time decay constant measured for step-stress relaxation experiments on protein condensates of increasing age. *p*-values derived from non-parametric Wilcoxon test. (**D**) Viscosity (*η = k · τ*) of the droplets as derived from the measurements in (C) and (D). Statistics derived from non-parametric Wilcoxon test. (**E**) Stiffness measured on the same protein condensates as in (**C**). Statistics derived from non-parametric Wilcoxon test.

**Fig. S4.**
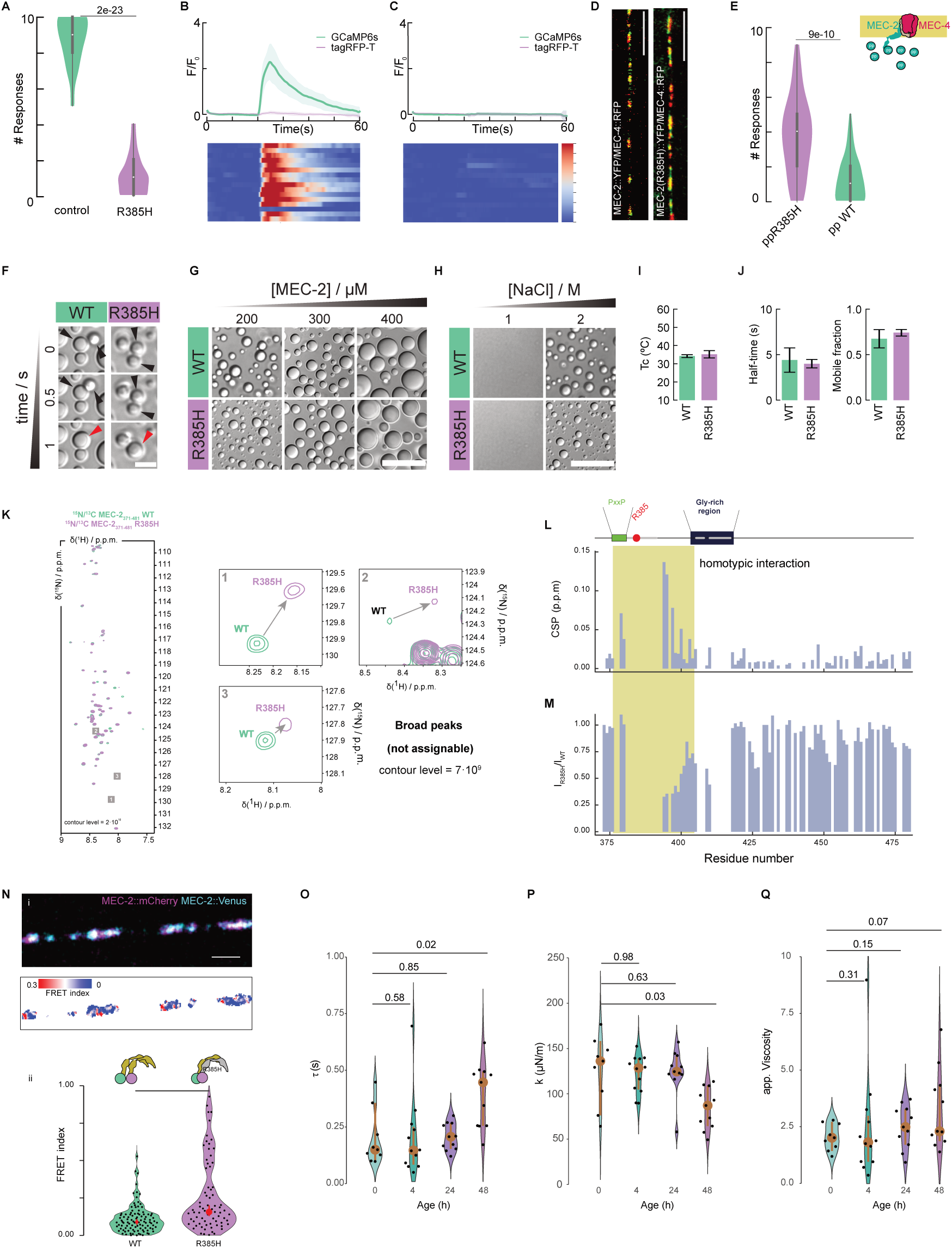
MEC-2 proline-rich domain is essential for the sense of touch. (**A**) Body touch response of wildtype vs. MEC-2(R385H) mutant. Vertical bar indicates SD, N≥ 60 animals. *p*-value derived from Kruskal-Wallis test. (**B** and **C**) Average GCaMP6s and calciumindependent tagRFP-T intensities recorded from TRNs of (**B**) wildtype and (**C**) MEC-2(R385H) mutant animals trapped inside the body wall chip. A 300kPa ’buzz’ stimulus was delivered for 2 s after recording 10 s baseline fluorescence. Individual recordings visualized as stacked kymographs (N=number of recordings). (**D**) Representative images of wildtype or MEC-2(R385H)::YFP mutant in green and MEC-4::RFP in red. Colocalization indicated in yellow. Scale bar = 10 µm (**E**) Touch response of wildtype animals with an overexpression of R385H mutant or wildtype proline-rich (PRiM) MEC-2 motifs specifically in TRNs. Mean±SD, N≥ 60 animals. *p*-value derived from Kruskal-Wallis test. Scheme of the experiment at the top right. (**F**) DIC microscopy images showing fusion events of 300 µM MEC-2 WT and R385H with 2 M NaCl at 37°C. Scale bar = 5 µm. (**G**) DIC microscopy images of MEC-2 WT and R385H at increasing protein concentrations of 200, 300 and 400 µM, with 2 M NaCl at 37 °C. Scale bar = 20 µm. (**H**) DIC microscopy images of 200 µM MEC-2 WT and R385H with 1 and 2 M NaCl at 37 °C. Scale bar = 20 µm. (**I**) T*_c_* value of the apparent absorbance measurement as a function of temperature of 200 µM MEC-2 WT and R385H mutant with 2 M NaCl. (**J**) Recovery half time and mobile fraction of MEC-2 WT and R385H mutant quantified from a FRAP experiment *in vitro* of a 370 µM sample with 2 M NaCl at 20 °C. (**K**) 2D ^1^H-^15^N NMR correlation spectra of MEC-2 WT and R385H mutation and the close-up of low-intensity (non-assignable) signals from 2D NMR spectra of the WT (green) and the R385H mutant (purple) MEC-2 C-terminal domain. (**L, M**) Chemical shift (CSP, **L**) and intensity ratio (**M**) for each residue for the comparison between WT and R385H. The lower intensities around the PRiM indicate a homotypic interaction between different MEC-2(R385H) molecules. (**N**) (i) Representative dual coloer color confocal image of the mixed MEC-2 (MEC-2::Venus and MEC-2::mCherry) population and the corresponding FRET map. Scalebar = 2µm. (ii) Distribution of FRET values derived from >50 measurements, showing a higher median (red dot) for the mutant MEC-2. (**O**) Time decay constant measured for step-stress relaxation experiments on protein condensates of increasing age, formed from MEC-2 (R385H) mutant. *p*-values derived from non-parametric Wilcoxon test. (**P**) Stiffness measured on the same protein condensates as in (**O**). Statistics derived from non-parametric Wilcoxon test. (**Q**) Viscosity (*η = k · τ*) of the droplets as derived from the measurements in (**O**) and (**P**). Statistics derived from non-parametric Wilcoxon test.

**Fig. S5.**
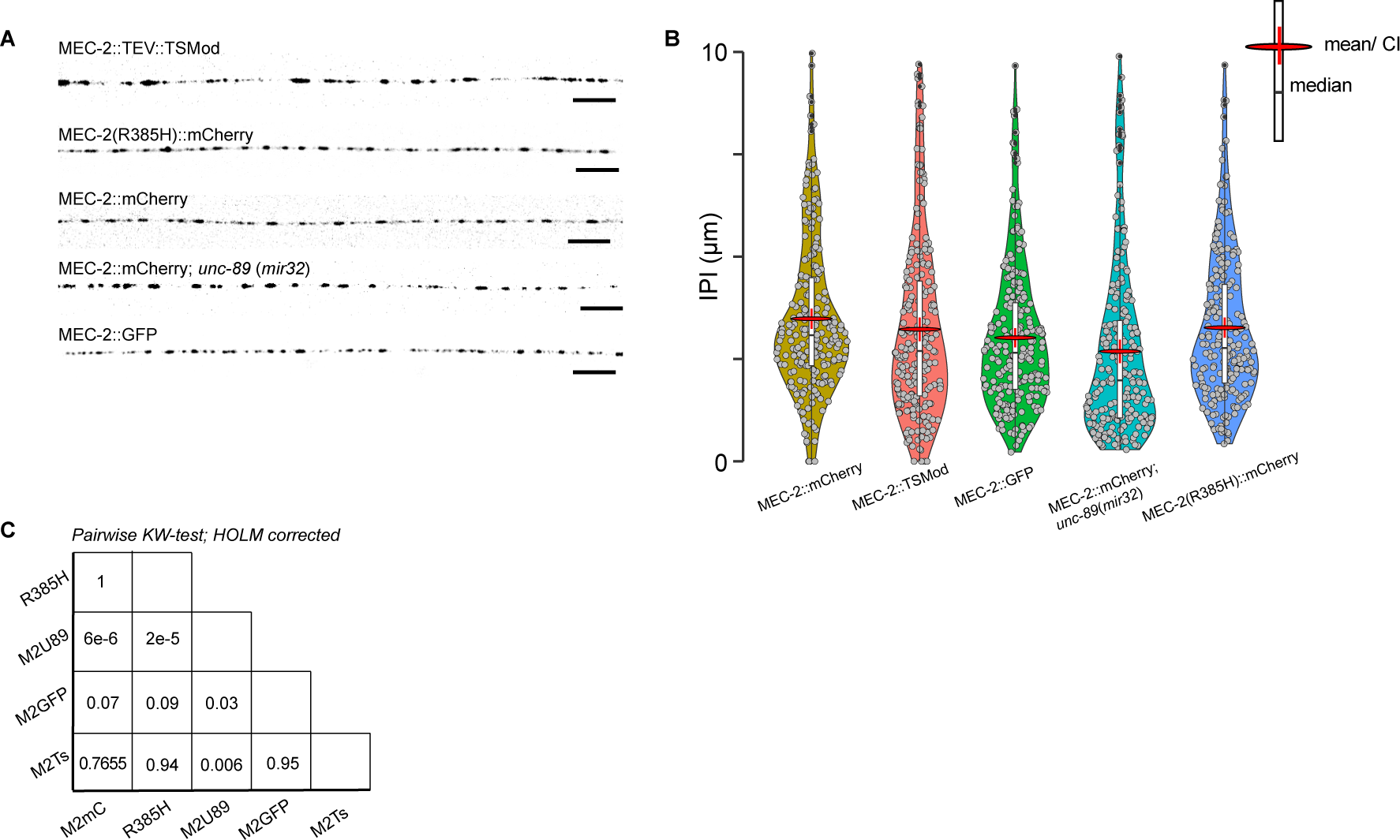
MEC-2 unchanged punctae pattern among genotypes. (**A**) Representative examples for the indicated MEC-2 alleles and genotypes. Scale bar = 10 µm. (**B** and **C**) Violin plot for all *mec-2* alleles and mutant backgrounds (**B**) and table indicating the *p*-values of the pairwise comparison (**C**) of their distribution from a Kruskall-Wallis test with Holm’s adjustment for multiple comparisons. The C-terminal truncated MEC-2 was not tested, as not interpunctum interval (IPI) could be extracted (IPI = 0 for a continuous distribution). Mean indicated as red lentil with vertical bar indicating the 95% confidence interval of the mean. Box encompasses 50% of all datapoint centered around the median (black, horizontal line). N = 230 MEC-2 puncta from 5-6 different animals’s TRNs.

**Fig. S6.**
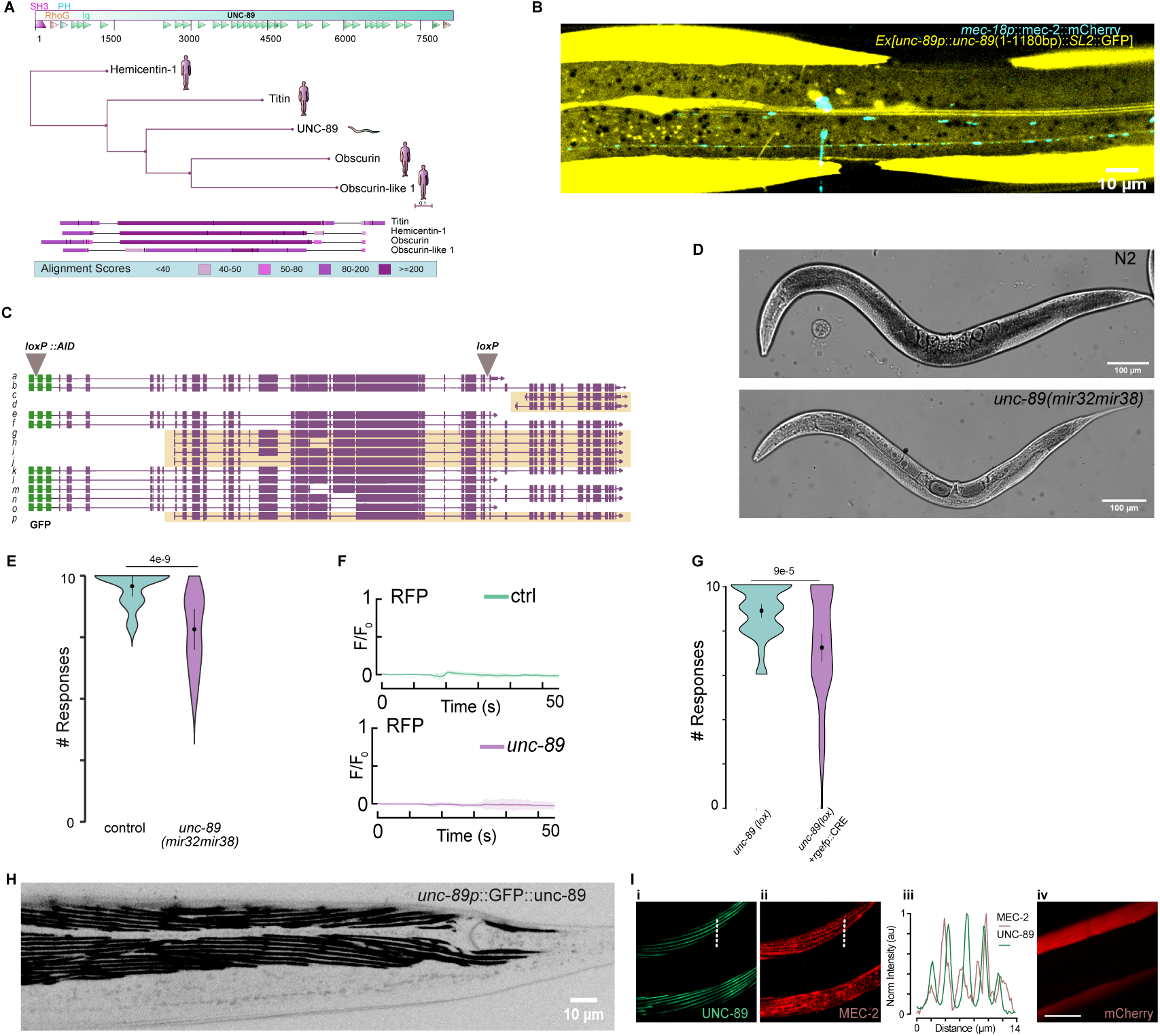
Non-muscular function of UNC-89. (**A**) Scheme of UNC-89 protein domains, BLAST against *Homo sapiens* genome and representation of the closest proteins Hemicentin-1, Titin, Obscurin and Obscurin-like 1 in a phylogenetic tree and protein alignment. (**B**) Representative micrograph of an animal expressing an *unc-89* promotor trap encompassing 4265 bp and the first 1180 bp of the genomic fragment (Exon1-Intron1-Exon2-Intron2-Exon3) showing expression of the largest isoforms in muscles and neurons. (**C**) Genomic organization of GFP-tagged *unc-89* locus and location of the two loxP sites. Yellow shadow shows remaining isoforms in the *unc-89(mir32)* allele knocking out the largest isoforms, which contains the SH3 domain. *mir32* was generated using a frameshift causing an abberrant initiation site. (**D**) Micrograph comparing young adult N2 and *unc-89(mir32)* animals. (**E**) Touch response of *unc-89(mir32)* KO allele compared to control wildtype animals. Circle indicates mean, vertical bar indicates 95% confidence interval, N=60 animals. *p*-value derived from Tukey HSD test. (**F**) Fluorescence intensity vs time of the calcium-independent fluorophore in the mechanical stimulation experiment showed in Fig. 3B. (**G**) Touch response of panneuronal (*rgefp*) knockout of UNC-89 compared to loxP flanked control animals in absence of CRE recombinase. Circle indicates mean, vertical bar indicates 95% confidence interval, N=60 animals. *p*-value derived from Tukey HSD test. (**H**) Representative fluorescence image of an animal with an N-terminal GFP tag at the endogenous locus of *unc-89* in frame with the SH3 domain. (**I**) Colocalization of MEC-2 heterologously expressed in body wall muscles with endogenous UNC-89 distribution (i, ii). (iii) Plot of the intensity profile taken on the dotted line indicated in i and ii. Soluble mCherry does not colocalize with UNC-89 (iv).

**Fig. S7.**
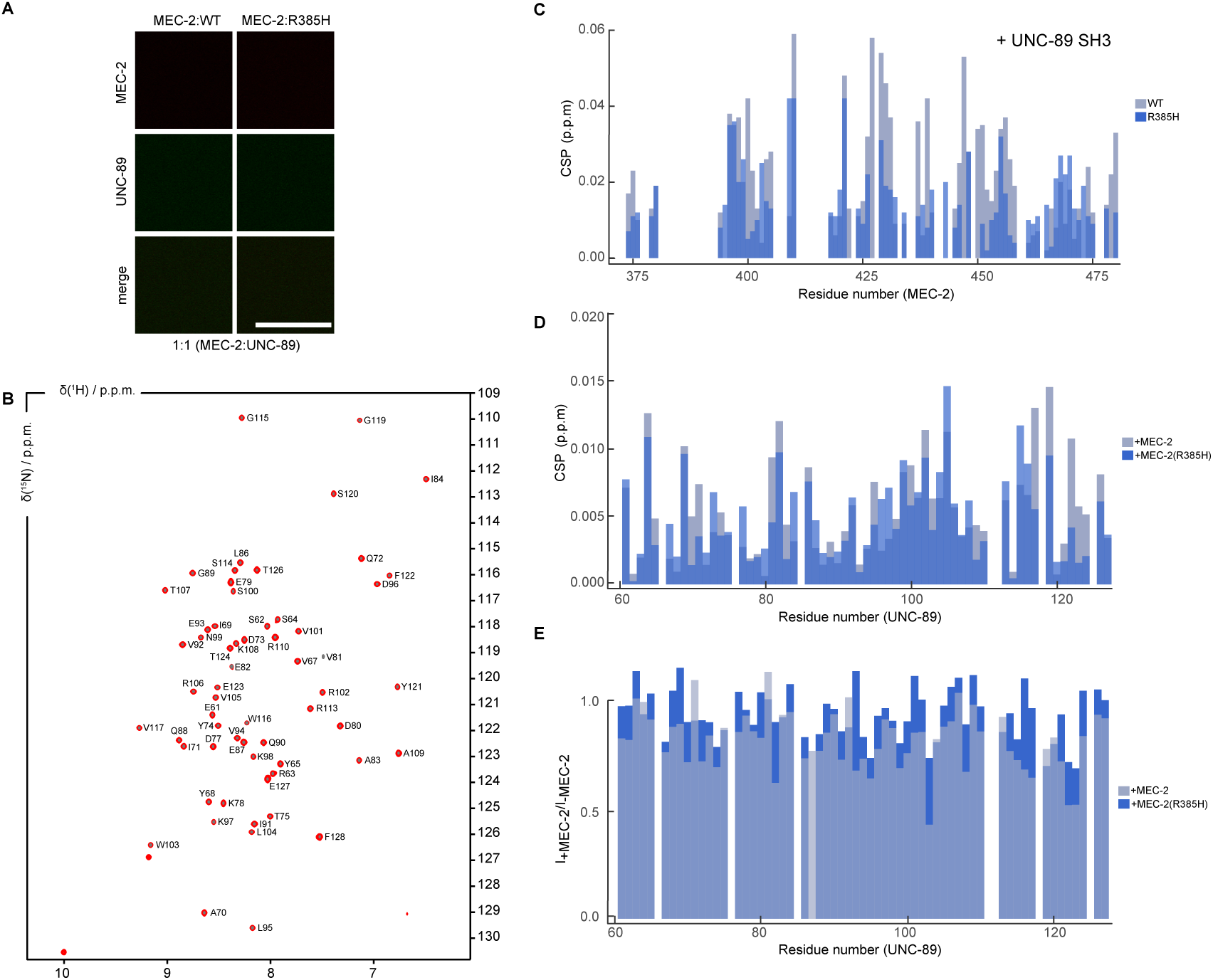
Structural changes upon binding of MEC-2 to UNC-89 SH3 domain. (**A**) Confocal fluorescence microscopy images of 200 µM MEC-2 C-terminus (WT or R385H mutant) labeled with Alexa Fluor 647 together with UNC-89 SH3 domain labeled with DyLight 488, at a molar ratio of 1:1 (MEC-2:UNC-89), with 2 M NaCl at 37 °C. Observed dissolution of MEC-2 droplets. Scale bar = 20 µm. (**B**) 61/62 H^N^ assignment annotated 2D NMR spectrum of the UNC-89 SH3 domain (BMRD ID: 51490). (**C**) Change in peak intensity of MEC-2 C-terminal wildtype (light blue) or R385H mutant (blue) upon binding to SH3 domain of UNC-89 (1:10 molar ratio). (**D**) Change in peak intensity of each of the SH3 residues of UNC-89 upon binding to wildtype (light blue) and R385H mutant (blue) MEC-2 (1:9 molar ratio). (**E**) Intensity ratio of the NMR spectra of the SH3 domain of UNC-89 in the presence or the absence of the C-terminal domain of MEC-2 (WT (light blue) and R385H mutant (blue)).

**Fig. S8.**
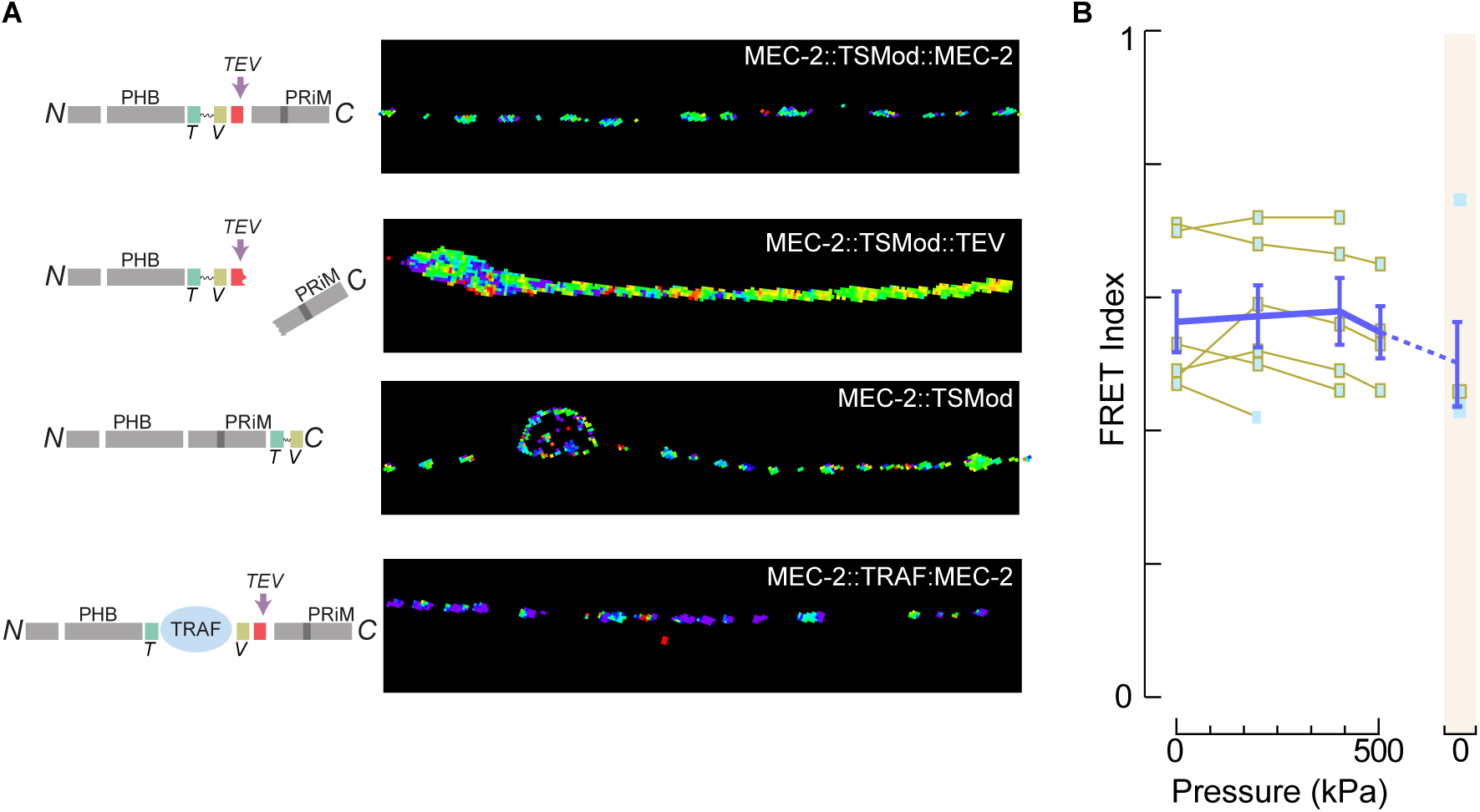
MEC-2::TSMod reports forces during body wall touch. (**A**) Scheme of the different tension sensor modules and control constructs employed, with their corresponding FRET maps. From top to bottom: TSMod in MEC-2 wildtype; in wildtype after TEV cleavage; in the C-terminal fusion as no force control; force-insensitive, low-FRET control replacing the flagelliform spring with a space domain domain. PHB, prohibitin domain; T, donor fluorophore; V, acceptor fluorophore; PRiM, proline-rich motif; TRAF, tumor necrosis factor receptor associated factor (*83*), TEV, Tobacco etch virus cleavage site. (**B**) FRET index changes with increasing pressure applied to the body wall of C-terminally tagged MEC-2 serving as a no-force control.

## Movies S1 to S5

**Movie S1. Organization and dynamics of MEC-2 in TRNs.** Representative videos of the MEC-2::mCherry dynamic and static pools in touch receptor neurons: 4 ALM and 4 PLM neurons.

**Movie S2. Dynamics of MEC-2 condensates *in vivo*.** Three representative videos of a MEC-2 condensate undergoing deformation (a), fission (b) and fusion (c) events during translocation along the neurite. Frame rate = 20 fps. Scalebar=2µm

**Movie S3. MEC-2 dynamics using FRAP.** Representative FRAP dynamics of the MEC-2::TEV::MEC-2 static pool and the C-terminally truncated construct after TEV coexpression.

**Movie S4. Fusion dynamics of MEC-2 condensates *in vitro*** Representative videos of the MEC-2 droplets undergoing fusion events *in vitro*.

**Movie S5. Touch-induced calcium transients.** Representative videos of a wildtype and a MEC-2::R385H mutant animal expressing a GCaMP6s calcium reporter in TRNs. A two second buzz was delivered after 10s. Top images are the calcium sensitive, lower images are the calcium insensitive channel (tagRFPt).

## Tables S1 to S5

**Table S1** List of candidate genes used for the RNAi experiment according to a preselection of C. elegans proteins with an SH3 domain (39) and proteins expressed in TRNs (40). Included the library chosen for each clone and the result of the touch assays after RNAi knock-down (Mean±SD).

**Table S2** List of strains used in this study.

**Table S3** List of plasmids and sequences used in this study.

**Table S4** List of CRISPR reagents: crRNAs and ssODNs used in this study.

**Table S5** List of touch response results for all the experiments, including Mean±SD and N for each assay.

